# Photoentrainment as Bayesian Inference: How the central circadian clock weights and integrate the intensity, wavelength and rate of change of natural light to estimate astronomical time

**DOI:** 10.64898/2026.10.27.762184

**Authors:** Danny M. Ball, Vincent Walsh, Russell G. Foster, Maarten Speekenbrink

## Abstract

Synchronising with the Earth’s 24-hour rotation using natural light as a timing cue presents a problem for the mammalian brain’s central circadian clock. Natural light is highly variable and can take a range of values at any given time of day, for example, fluctuating due to differences in weather conditions and across seasons.

Despite this variability, the central clock maintains close alignment with astronomical time, and here we describe how this may be achieved computationally. We present a Bayesian model in which the clock’s endogenous oscillation is used to predict the probability of observing the features of light, intensity, spectral composition, and their co-occurrence given the time of day, drawn from a distribution of values typically experienced. When observed light falls outside the predicted range, a prediction error provides evidence that the clock may be misaligned with astronomical time. How much the clock then adjusts its phase depends on the relative uncertainty associated with its internal prediction and observed light. These are combined as a weighted average according to their relative reliability, drawing the clock’s updated phase towards whichever provides the more reliable estimate, either the clock’s prediction or the observation, of astronomical time. Using natural light measurements collected over one week from 59 human observers and 100 days of environmental spectral measurements, we characterise how the variability of natural light systematically changes across the day and show how the clock may exploit this structure for circadian timekeeping. The model explains how the canonical shape of the phase-response curve emerges from the relationship between light distributions and time of day, and how properties of light, including intensity and spectral composition, can be integrated according to their reliability. The central clock’s synchrony to light thus represents a specific instance of a general computational strategy employed by the mammalian nervous system, whereby sensory information is interpreted relative to an internally generated prediction.

## Introduction

A fundamental challenge for the mammalian central circadian clock is to extract biologically meaningful timing information from a noisy and perpetually fluctuating light environment^1–3^. The central clock, the suprachiasmatic nucleus (SCN), is the mammalian master pacemaker, synchronising the timing of the daily cycles of physiology and behaviour, collectively known as circadian rhythms^4–6^. While the central clock has an endogenous oscillation close to 24 hours, it does not keep perfect time and displays day-to-day variability in its circadian period, i.e. the length of one subjective day^7–9^. Therefore, the central clock requires daily adjustments so that subjective circadian time is aligned with astronomical time, which is determined by the Earths 24-hour rotation. The light-dark cycle is a close proxy for astronomical time. The solar angle, the angular position of the Sun relative to the observer, changes throughout the day as the Earth rotates, thereby altering both the quantity (intensity) and the quality (spectral composition) of light, and is the primary timing cue for the circadian system in mammals^10,11^. Through a process called photoentrainment, changes in ambient light are captured by specialised ganglion cells in the retina, which express the photopigment melanopsin with a response sensitivity peak to short-wavelength light (480 nm). Other retinal cells contribute to photoentrainment, cones for detecting the spectral changes in light and rods for changes in dim light are integrated at the level of ganglion cells, which transmit to the central clock, advancing or delaying its phase if it is running slow or fast relative to astronomical time^12–15^.

However, photoentrainment in natural light presents a problem for the circadian clock. For example, while the intensity of natural light spans many orders of magnitude, providing a reliable difference between day and night, light can have significantly different values at the same time across days. For example, light intensity at 9 am one day can differ significantly from 9 am the next due to differences in weather conditions and across seasons. Moreover, different times of the same day can have similar intensity values. A dim midday under cloud cover and a clear evening can be equally bright, causing ambiguity about solar elevation^10^. To reduce ambiguity about the time of day, the central clock integrates multiple features of light^1,10,11,16^. This is especially informative at twilight, during the transition of dawn and dusk. As the sun approaches and passes below the horizon, direct sunlight is lost and the remaining light is dominated by scattered skylight^17^, such that ambient light contains a higher proportion of short wavelengths than daylight^10,16^. The familiar redness of the setting sun is the same process seen from the other side: short wavelengths are scattered out of the direct beam, which is why it reddens, and it is that scattered light which dominates the sky once the sun is below the horizon. The duration of light exposure and the direction and rate of change of intensity and spectral composition also carry time information. However, a problem arises when the information provided by these components does not agree. Any individual feature of light may be intermittent or change rapidly due to behaviour, for instance the flickering of light while passing under trees, or changes in spectral composition as light reflects off different surfaces, which may provide conflicting information about the time of day. It is not understood how the central clock resolves this potential conflict between timing signals or how much it should adjust its phase, given that any cue can typically arise from a range of possible values. The purpose of this manuscript is to describe how the central clock maintains synchrony with the solar day under such uncertainty.

Previous mathematical models of the central clock have successfully described circadian oscillators and phase resetting to light, accurately reproducing many experimental observations^18–22^. Where variability has been introduced into such models, it has generally been treated as something the clock needs to reduce or remove^23^. However, it is feasible that the variability in natural light may actually be useful for circadian timekeeping. Light does not just vary. It varies systematically across the day (Figure 1). For example, light during the middle of the day is not only brighter than at midnight, but also substantially more variable, meaning that the range of light intensities typically experienced is greater than the relatively constrained range observed at night^24^. This may help the central clock weight timing cues according to their reliability, that is, according to how probable they are at different times of day, and combine them to estimate astronomical time. This aligns with a Bayesian perspective, which suggests that brains represent the world probabilistically, using quantities such as central tendency and uncertainty to interpret sensory input, with variability itself forming part of the computational process^25–28^. Using this logic, circadian photoentrainment can be understood as a form of Bayesian inference^29^, in which the clock uses its internal phase (the prior; the time represented by the clock) and the degree of certainty about it to generate predictions about how likely it is to observe a given timing cue (the likelihood), for instance, the decrease in intensity and shift in spectral composition of ambient light in the evening. These predictions are combined with incoming sensory evidence in a weighted average to arrive at the most probable estimate of true astronomical time (the posterior). We show how this framework reproduces the shape of the circadian phase response curve^30^. Because bright light is far more probable during the day than at night, the same light exposure produces larger phase shifts when it occurs at unexpected times. We further show how the clock integrates the intensity and wavelength of light as complementary sources of evidence about solar time, weighting each according to how reliably it predicts the time of day under prevailing conditions. We demonstrate how the direction and rate of change of these features disambiguate dawn from dusk, resolving an ambiguity that intensity and wavelength alone cannot. Thus, circadian photoentrainment may be a specific instance of a general computational strategy found throughout the mammalian nervous system, in which perception is relative and is evaluated compared to an internally maintained prediction^31–33^.

**Figure 1:**
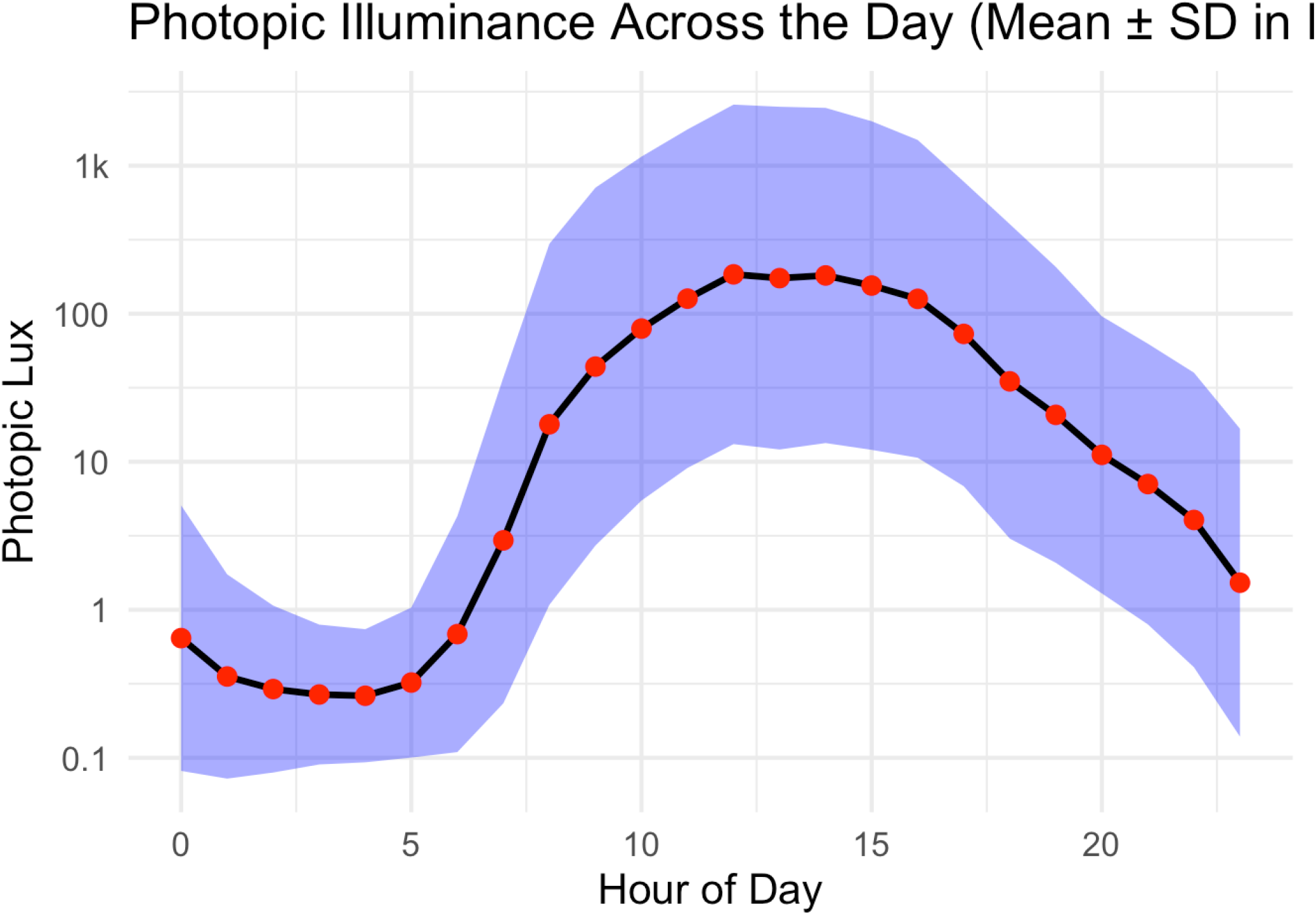
photopic illuminance magnitude and variability across the 24-hour day. Photopic illuminance was measured for one week in 59 UK adults using a wrist-worn spectrometer, with data taken from Didikoglu et al.^24^. The x-axis represents clock time, divided into 1-hour intervals, and the y-axis shows photopic illuminance on a log_10_ (lux) scale. Light exposure measurements were pooled across participants and averaged within each hour of the day. Photopic illuminance values were log_10_-transformed because environmental light exposure spans several orders of magnitude and is strongly right-skewed. For each hour, the mean and standard deviation were calculated in log space. The black line and red points show the hourly mean, while the shaded region indicates ±1 standard deviation. The plot shows higher light levels and greater variability during the day, consistent with changing exposure as people move between indoor and outdoor environments, as well as natural variability in ambient light possibly due to factors such as cloud cover. At night, light levels are lower and less variable, reflecting more stable dim light exposure.

## Bayesian Observer Model of the Central Circadian Clock

We model the central circadian clock as a Bayesian observer that infers astronomical time from multiple features of the natural light environment. Astronomical time is treated as a latent state, a quantity that must be estimated from sensory evidence. The clock’s endogenous rhythm provides an ongoing prediction of this latent state, continuously updated as new photic evidence arrives, with the magnitude of each update determined by how informative the current observation is relative to the clock’s existing prediction. The principal components are described below (Figure 2).

**Figure 2:**
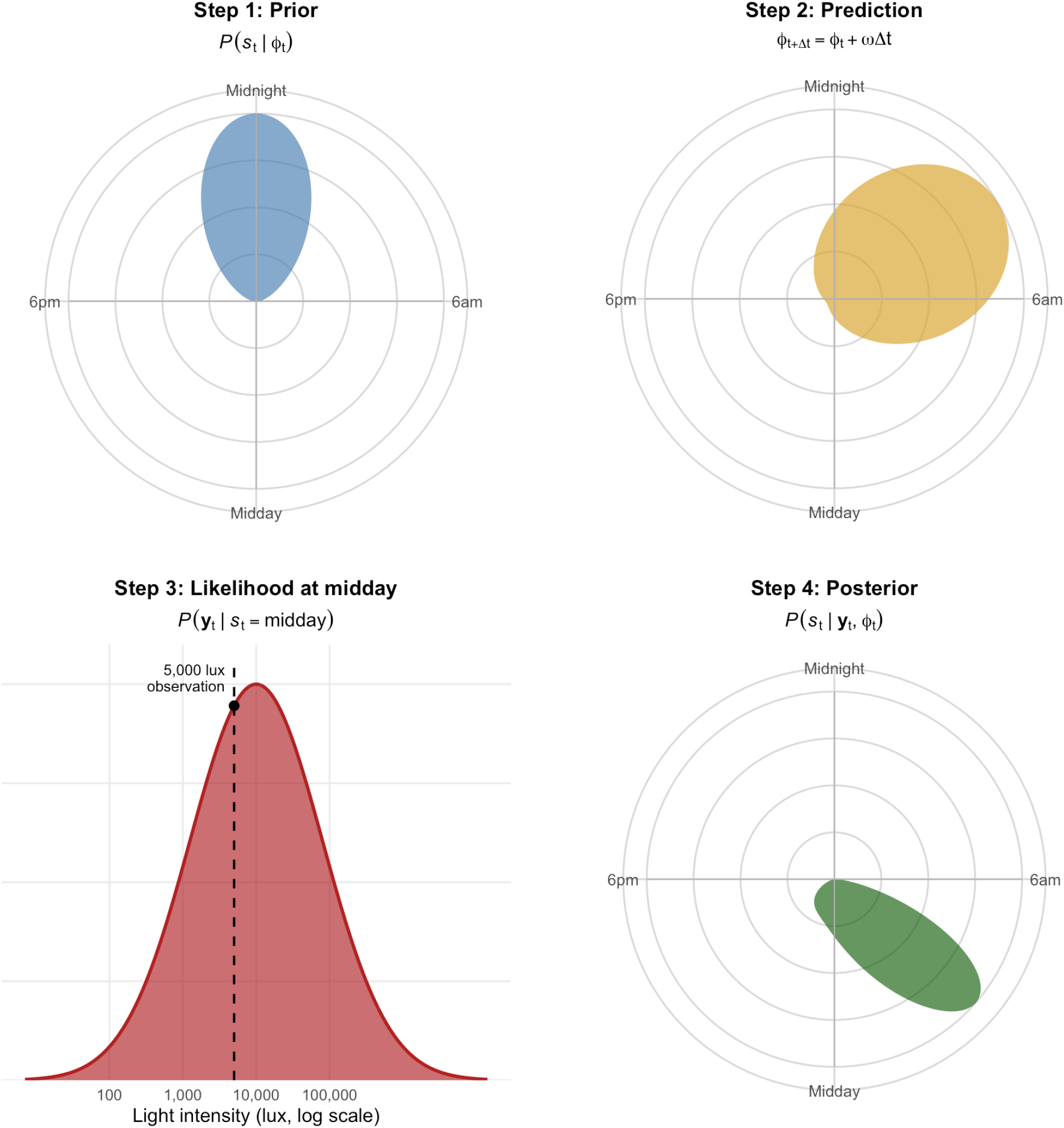
Bayesian Clock Model. The Bayesian clock model provides a framework for understanding how the circadian central clock dynamically estimates and adjusts its internal representation of time. In this model, the central clock’s estimate of time is represented as a probability distribution over a continuous 24-hour circle, consistent with the treatment of astronomical time as a continuous latent variable in the formal model. The process begins with the internal clock’s prior belief about the current time (Step 1, top left panel), distributed around the circle. Initially, the belief is centred around the most probable time, in this figure midnight, with confidence gradually decreasing for times further away. This distribution reflects the clock’s uncertainty about its phase in relation to the astronomical day. As time progresses, the clock propagates its belief forward around the circle to account for the natural advancement of the circadian cycle in the absence of external cues (Step 2, top right panel). This prediction step reflects the passage of time, shifting the belief distribution forward and broadening it, since the clock’s internal timekeeping is not perfectly precise and uncertainty accumulates during periods when no light or environmental information is available, such as during sleep. When the clock receives a light cue, here a light intensity of 5,000 lux, it evaluates how well this observation matches the light typically expected at each candidate hour of the day (Step 3, bottom left panel). This observation is closest to the mean of the light distribution typically observed around midday, producing a likelihood that peaks sharply at midday and falls away toward midnight. Finally, the clock combines this likelihood with its propagated prior belief to produce an updated, posterior estimate of the current time (Step 4, bottom right panel). Because the observation is closest to the midday distribution, the posterior is drawn toward midday, reflecting a substantial phase adjustment away from the clock’s propagated estimate of the early hours of the morning. More generally, stronger and more surprising light cues of this kind produce a greater adjustment of the clock’s internal estimate, while weaker or expected cues allow the prior belief to dominate. This posterior belief then becomes the new prior for subsequent cycles, as the clock continuously updates its estimate as new evidence arrives. In the circular panels (Steps 1, 2 and 4), the angle is the time of day (midnight at the top, 6am on the right, midday at the bottom and 6pm on the left) and the distance from the centre is the probability assigned to that time, scaled to the same peak height. In Step 3, the x-axis is light intensity (lux, log scale) and the y-axis is probability density, scaled to the same peak height.

### Latent state

Astronomical time is represented as a circular variable spanning one full solar day:

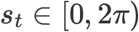

where *s*_*t*_ denotes astronomical time at time *t*, with 2*π* corresponding to 24 hours.

### The central clock’s endogenous rhythm

The clock’s phase advances continuously:

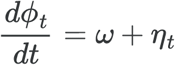

where *ϕ*_*t*_ is the circadian phase at time *t, ω* = 2*π*/*τ* is the endogenous angular frequency determined by the individuals intrinsic circadian period *τ*, and *η*_*t*_ represents random fluctuations in the rate of phase progression. Each fluctuation is independent of the last, but their effects accumulate, so that uncertainty about phase grows over time.

### Photic observations

The clock receives multiple features of the light environment that vary systematically with time of day:

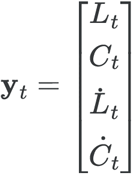

where *L*_*t*_ is log-transformed light intensity, *C*_*t*_ is spectral composition, and 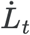 and *Ċ*_*t*_ are their rates of change, estimated over a short integration window *δ*:

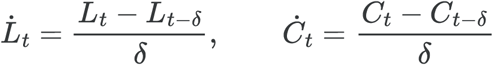

Intensity and spectral composition change systematically with the elevation of the sun, so the same pair of values (*L*_*t*_, *C*_*t*_) can occur on either side of solar noon. The rates of change resolve this. Their sign, whether light is getting brighter or dimmer and shifting toward or away from short wavelengths, indicates which half of the day the observer is in, while their magnitude, how quickly this is happening, provides a further, independent cue.

### Prior distribution

The clock’s belief about astronomical time is represented as a von Mises distribution centred on its current circadian phase:

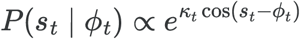

We use the von Mises distribution, the circular analogue of the Gaussian, because *s*_*t*_ is a circular quantity: a distribution that did not wrap around the day boundary would treat 23:59 and 00:01 as far apart, when for a clock they are one minute apart. The von Mises distribution avoids this, ensuring that estimates and updates behave consistently regardless of where on the 24-hour cycle they occur.

*κ*_*t*_ is the concentration parameter. Higher values of *κ*_*t*_ correspond to a narrower, more confident prior; lower values to a broader distribution and greater uncertainty. *κ*_*t*_ follows from the model itself: it increases when photic evidence sharpens the posterior and decreases between updates as phase uncertainty accumulates.

### Prediction

Between updates, the clock’s belief is carried forward by its endogenous rhythm. Over an interval Δ*t*, its centre advances at the endogenous rate:

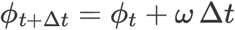

and its concentration *κ*_*t*_ decreases as phase uncertainty accumulates. In the absence of light, the clock’s estimate therefore continues to advance while becoming progressively less certain.

### Likelihood

The likelihood describes the probability density of the observed light features given each candidate time of day, and is estimated empirically from the statistical structure of natural light. For light intensity alone:

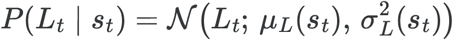

where *μ*_*L*_(*s*_*t*_) and 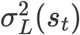 are the mean and variance of log intensity typically observed at time *s*_*t*_. Because intensity enters in log units, this corresponds to a log-normal distribution of intensity in lux. For intensity and spectral composition together:

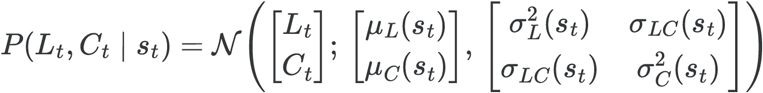

where *μ*_*C*_(*s*_*t*_) and 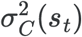 are the mean and variance of spectral composition at time *s*_*t*_, and *σ*_*LC*_(*s*_*t*_) is the covariance between intensity and spectral composition. Because this covariance changes with time of day, the joint pattern of the two cues can carry information about astronomical time beyond that carried by either cue alone. More generally, for the full set of features:

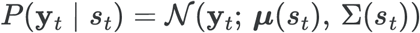

where ***μ***(*s*_*t*_) and Σ(*s*_*t*_) are the mean vector and covariance matrix of the light features typically observed at time *s*_*t*_.

### Posterior distribution

At each small step Δ*t*, the clock combines its prior with the likelihood of the observed light using Bayes rule:

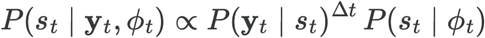

The clock updates in small discrete steps, and the evidence from each step is weighted by its duration Δ*t* (in hours), so that sustained light contributes more evidence than brief light and the result is essentially independent of how finely time is divided. Because the likelihood, viewed as a function of *s*_*t*_, is not von Mises in form, the posterior is not von Mises either. It is therefore approximated by the von Mises distribution with the same circular mean and mean resultant length, which gives the updated phase 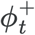 and concentration 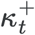. These then form the prior for the next step. The change in the clock’s estimate at each step is 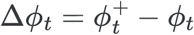, wrapped to (−*π, π*] and expressed in hours as Δ*ϕ*_*t*_ × 24/2*π*, with positive values indicating phase advances and negative values phase delays. The phase shift produced by a light pulse is the sum of these changes over its duration.

The difference between the clock’s prediction and the observed light constitutes a prediction error. The larger this error, and the less certain the clock is of its prediction, the greater the update. Because the posterior at one moment becomes the prior for the next, evidence accumulates recursively over time. This gives the clock a time-of-day-dependent sensitivity to light: during the day, when bright light is expected and naturally variable, the clock is largely unaffected by brief fluctuations in light, since a given observation typically produces only a small prediction error; at night, when light is not expected at all, the same observation produces a large prediction error and a correspondingly large update. The same recursion also accounts for the saturation effect, whereby the same light produces progressively smaller phase shifts the longer it continues: as consistent evidence accumulates, *κ*_*t*_ increases and the posterior becomes more concentrated, so the clock grows increasingly certain of its estimate and each further update produces a progressively smaller shift.

## A Bayesian Description of the Phase Response Curve to Light

A phase response curve (PRC) to light, (Figure 3) describes how the direction and magnitude of circadian phase shifts change in magnitude as a function of the time of the subjective day at which light is received. Light during the subjective morning advances the clock, light during the subjective evening and early night delays it, and light during the subjective day produces little or no shift, a region of relative insensitivity often called the dead zone^30^. The size of the shift is largest around subjective midnight and diminishes toward midday producing the characteristic shape of the PRC.

**Figure 3:**
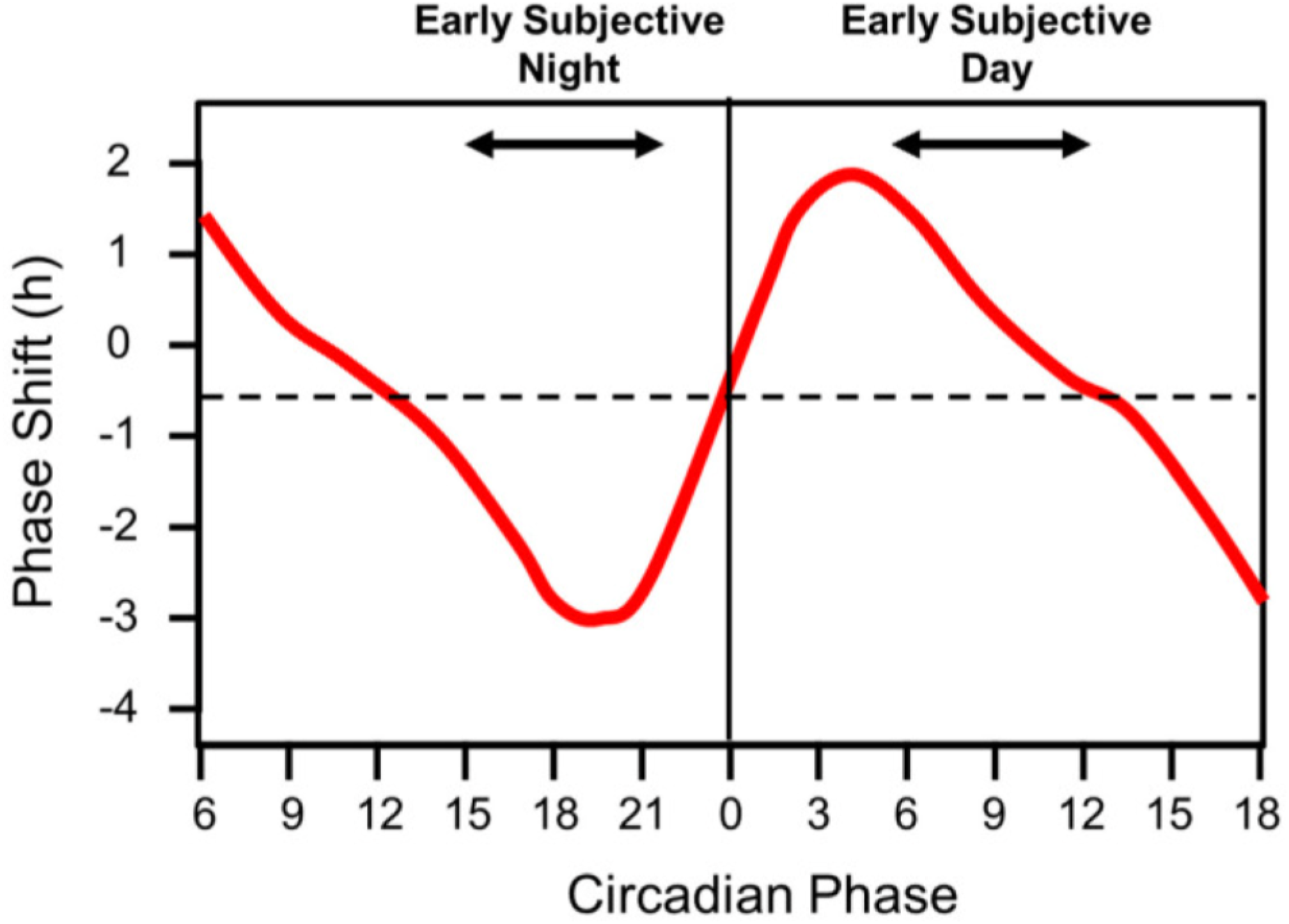
Phase Response Curve to Light: In a Phase response curve, the x-axis represents subjective circadian time (CT), the phase of the central circadian clock. The y-axis represents the magnitude of a phase shift in response to a light pulse. A value of zero on the y-axis corresponds to no phase advance or delay, while positive values indicate phase advances and negative values indicate phase delays. Light exposure during the early circadian night typically produces phase delays, whereas light exposure during the late circadian night or early circadian morning produces phase advances. During the circadian day, light generally produces little or no phase shift, creating a region of relative insensitivity often referred to as the “dead zone”. The PRC provides a framework for understanding how environmental light synchronises the internal clock to the 24-hour day and helps explain entrainment. The shape and amplitude of the PRC may vary across individuals and depend on factors including the intensity and wavelength of light, the duration of the pulse and prior light history. Adapted from Foster et al.^1^.

The characteristic shape of the PRC can arise from the Bayesian model’s response to the statistical structure of natural light, i.e., the probability of encountering a given light intensity at different times of the astronomical day. To illustrate this, we calculated the probability of observing 5,000 lux at midday and midnight (Figure 4). 5,000 lux was selected because it has been shown to reliably elicit phase advances and delays in humans^34^, using group-averaged photopic illuminance measured over one week in 59 UK adults wearing a wrist-worn light sensor^24^. We fit probability density functions to capture the range of light intensities averaged across the population. We then calculated how likely it would be to experience a light of 5,000 lux. The probability of observing 5,000 lux was 0.028 at midday and 6.9 × 10^−6^ at midnight, making 5,000 lux approximately 4,000 times more probable during the day than at night. Rather than responding directly to the physical magnitude of a light stimulus, the Bayesian observer evaluates how probable the observed light level is under the learned statistical structure of the environment. The same intensity therefore carries different information depending on when it occurs: a level that is common during the day but rare at night provides strong evidence that the clock is misaligned.

**Figure 4:**
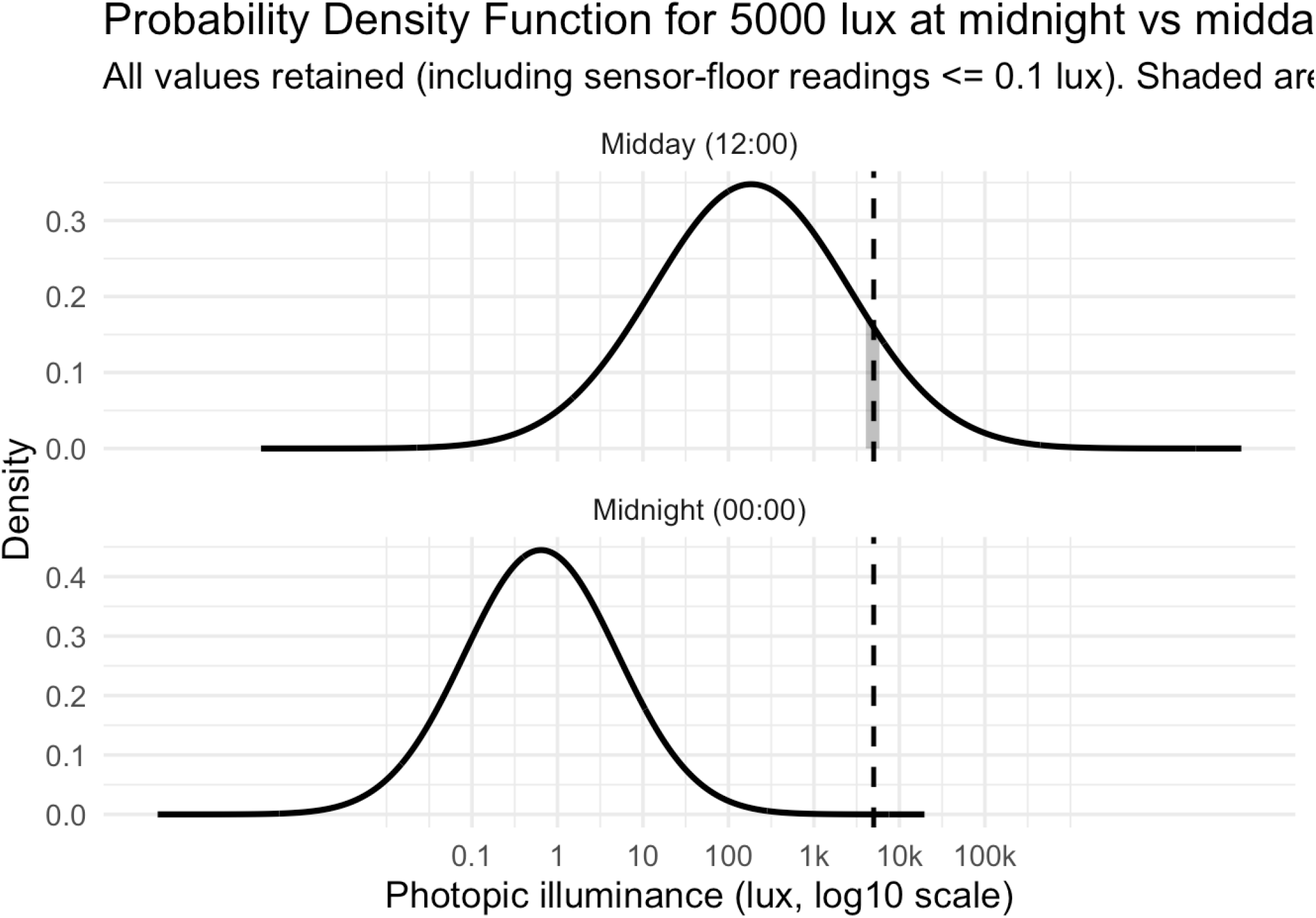
Probability distributions of log photopic light exposure of 5000 lux at midnight and midday. Probability density functions show Gaussian fits to log-transformed photopic illuminance at midnight (00:00) and midday (12:00). The x-axis shows photopic illuminance on a log_10_(lux) scale, and the y-axis shows probability density. At midday, the fitted distribution has a mean of 2.32 log_10_(lux) with a standard deviation of 1.02 log units, corresponding to a typical range of approximately 20 to 2,175 lux (±1 SD), although empirical daytime exposures extend much higher, reaching approximately 58,000 lux at the 99th percentile and >145,000 lux at maximum. At midnight, the fitted distribution has a mean of 1.00 log_10_(lux) with a standard deviation of 0.83 log units, corresponding to a typical range of approximately 1.5 to 68 lux (±1 SD), with most observations occurring at low light levels. A vertical dashed line indicates an example light level (5000 lux; log_10_ ≈ 3.7), which lies within the daytime range but falls in the extreme upper tail of the nighttime distribution.

Applying this principle at each hour of circadian phase generates a model-derived phase response curve (Figure 5). A one-hour pulse of 5,000 lux was applied at each internal circadian hour, with one hour of light counted as one hour of evidence, as throughout, and the resulting phase shift was calculated by updating a von Mises prior centred on that hour with a log-normal likelihood derived from the empirical hourly light distributions. The difference between the clock’s prediction and the observed light level constitutes a prediction error: the larger this error, the greater the update to the clock’s estimate of astronomical time.

**Figure 5:**
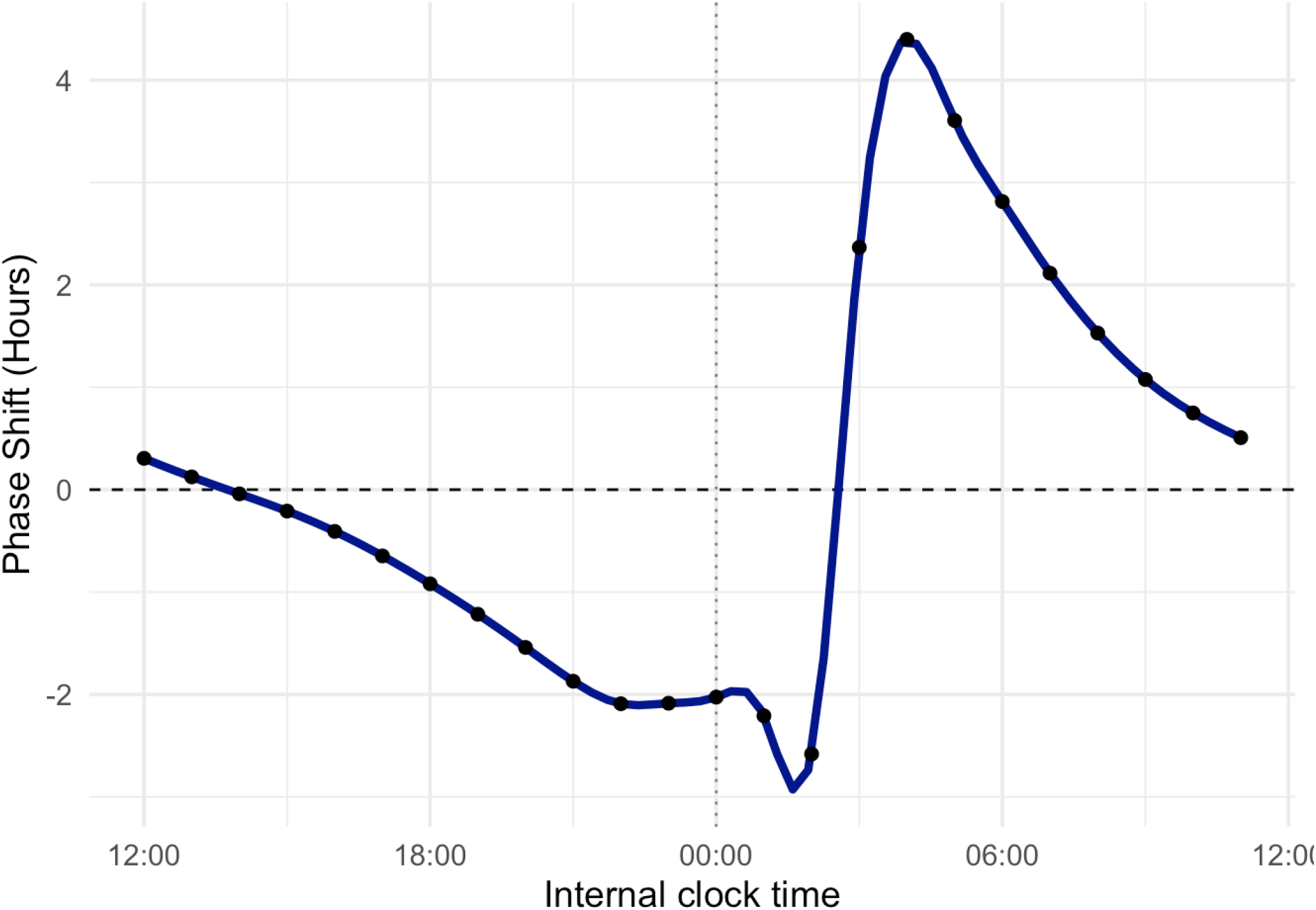
A phase response curve was generated by the Bayesian observer model in response to a one-hour pulse of 5,000 lux applied at each internal circadian hour. For each hour, a prior belief about circadian phase was represented as a von Mises distribution centred on that hour, and a likelihood was calculated from the observed light intensity (log-transformed) relative to the hourly mean and variance of empirical photopic illuminance. The prior and likelihood were combined to form a posterior belief about circadian phase, and the phase shift was calculated as the circular difference between the mean of the prior and the mean of the posterior. The hour’s evidence is applied in a single update rather than accumulated across the hour, so the within-pulse saturation shown in Figure 15 is not modelled here. The x-axis represents internal circadian time (hours), centred on midnight, and the y-axis shows the resulting phase shift in hours, with positive values indicating phase advances and negative values indicating phase delays. The curve produced by the model reflects the differential probability of observing 5,000 lux across the circadian cycle: because this intensity is highly improbable during the subjective night, light exposure at these hours generates a large mismatch between predicted and observed light, and therefore a large phase shift, whereas exposure during the subjective day, when 5,000 lux is common, produces comparatively little shift.

Because 5,000 lux is highly improbable during the subjective night, light exposures at these times generate large prediction errors and correspondingly large phase shifts. During the subjective day, the same light falls within the expected range, prediction errors are small, and phase adjustments are correspondingly modest. The model produced the characteristic shape of a human phase response curve, with the largest phase advance of approximately 5.8 hours at 03:00 and the largest phase delay of approximately 3.8 hours at 02:00. The same approach was extended across light levels of 1, 10, 100, 1,000, and 10,000 lux (Figure 6). At low intensities (1 and 10 lux), phase shifts during the day were small but opposite in direction to those produced by brighter light, whereas at higher intensities, progressively larger advances and delays emerged during the biological night and early morning. This asymmetry follows from the Bayesian framing of the model: the clock does not respond to light as a fixed physical stimulus, but interprets it as evidence about whether the clock is correctly aligned with astronomical time. A low-light pulse during the day is unexpected, since daytime light is typically bright, and therefore signals misalignment, whereas the same low-light pulse at night is unsurprising and carries comparatively little information. Brighter light shows the reverse pattern: it is uninformative during the day but highly informative at night, since bright light is rare during the biological night. Larger phase shifts therefore arise not simply because a stimulus is more intense, but because its occurrence is more improbable given the expected light level at that circadian phase.

**Figure 6:**
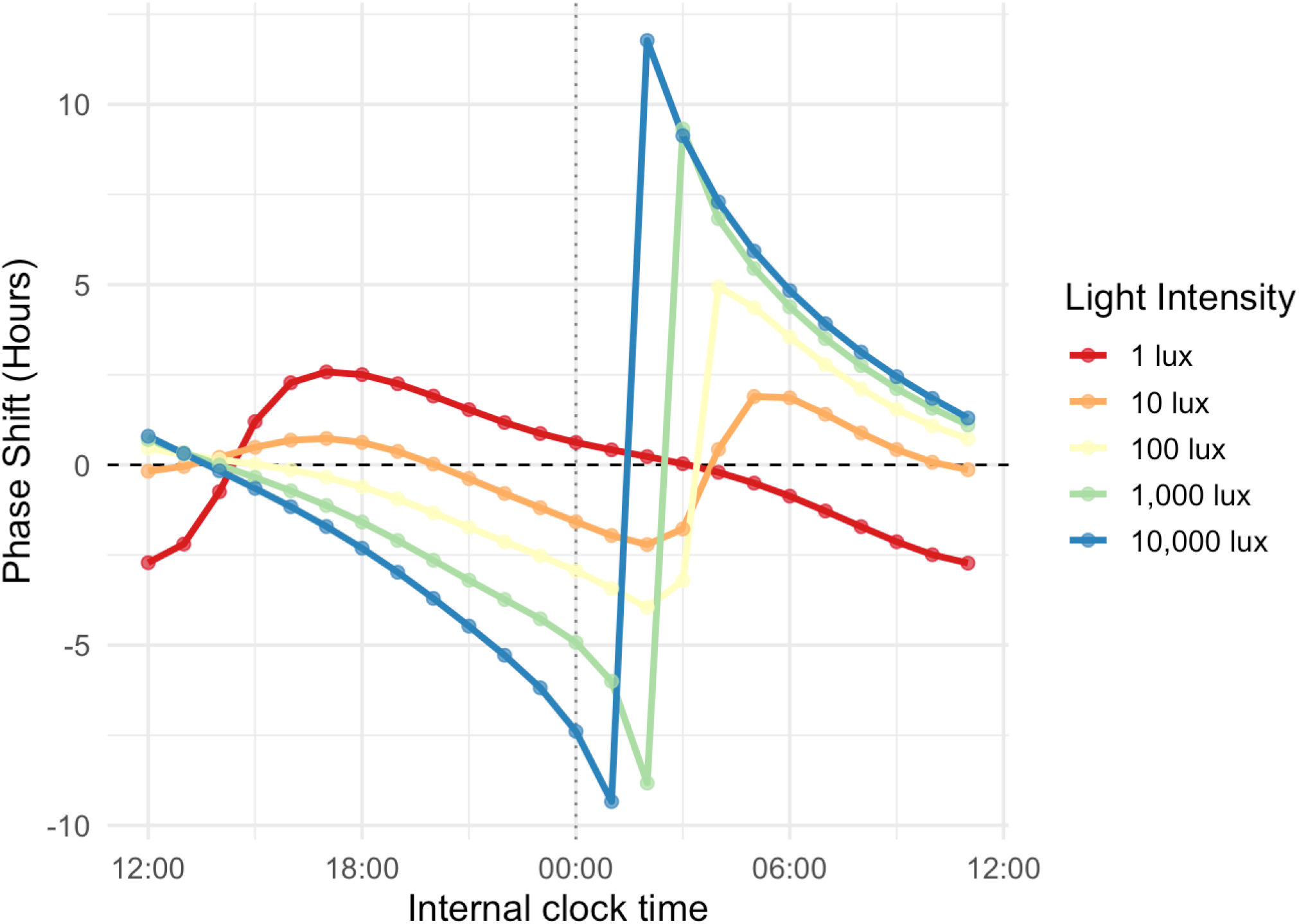
Phase response curves generated by the Bayesian observer model for one-hour pulses across multiple light intensities (1, 10, 100, 1,000, and 10,000 lux). The x-axis represents internal circadian time (hours), centred on midnight, and the y-axis shows phase shift (hours), where positive values indicate phase advances and negative values indicate phase delays, and zero indicates no change in phase. Phase shifts were computed by combining a fixed prior belief with a likelihood estimated from the hourly mean and variance of log-transformed photopic illuminance. As light intensity increases, the resulting phase shifts become progressively larger, because brighter light is less probable at a given circadian phase and therefore provides stronger evidence of misalignment. At the lowest intensities (1 and 10 lux), phase shifts during the day were small but opposite in direction to those produced by brighter light, reflecting the fact that low light is itself unexpected during the biological day.

## Individual Differences in Light Sensitivity to the Same Light Stimulus

People differ in how much their circadian clock responds to the same light stimulus. In our model, this can arise from two of its components, the likelihood and the prior. Because the likelihood is derived from a persons own light history, the model predicts that people will differ in their phase response to the same light. To illustrate this, PRCs were generated for each participant using likelihoods estimated from their own hourly light exposure, with the stimulus fixed at a one-hour pulse of 5,000 lux (Figure 7). Despite the identical stimulus, the resulting PRCs varied systematically across participants. Most showed the characteristic shape, but the timing of peak sensitivity differed, reflecting when bright light typically occurs in each person’s day. People with brighter or later evening light exposure assign a higher probability to 5,000 lux in the evening, reducing the prediction error at those times and shifting the phase of maximum sensitivity later into the night. The amplitude of the PRC also differed, because the probability of encountering 5,000 lux in the evening differed between participants.

**Figure 7:**
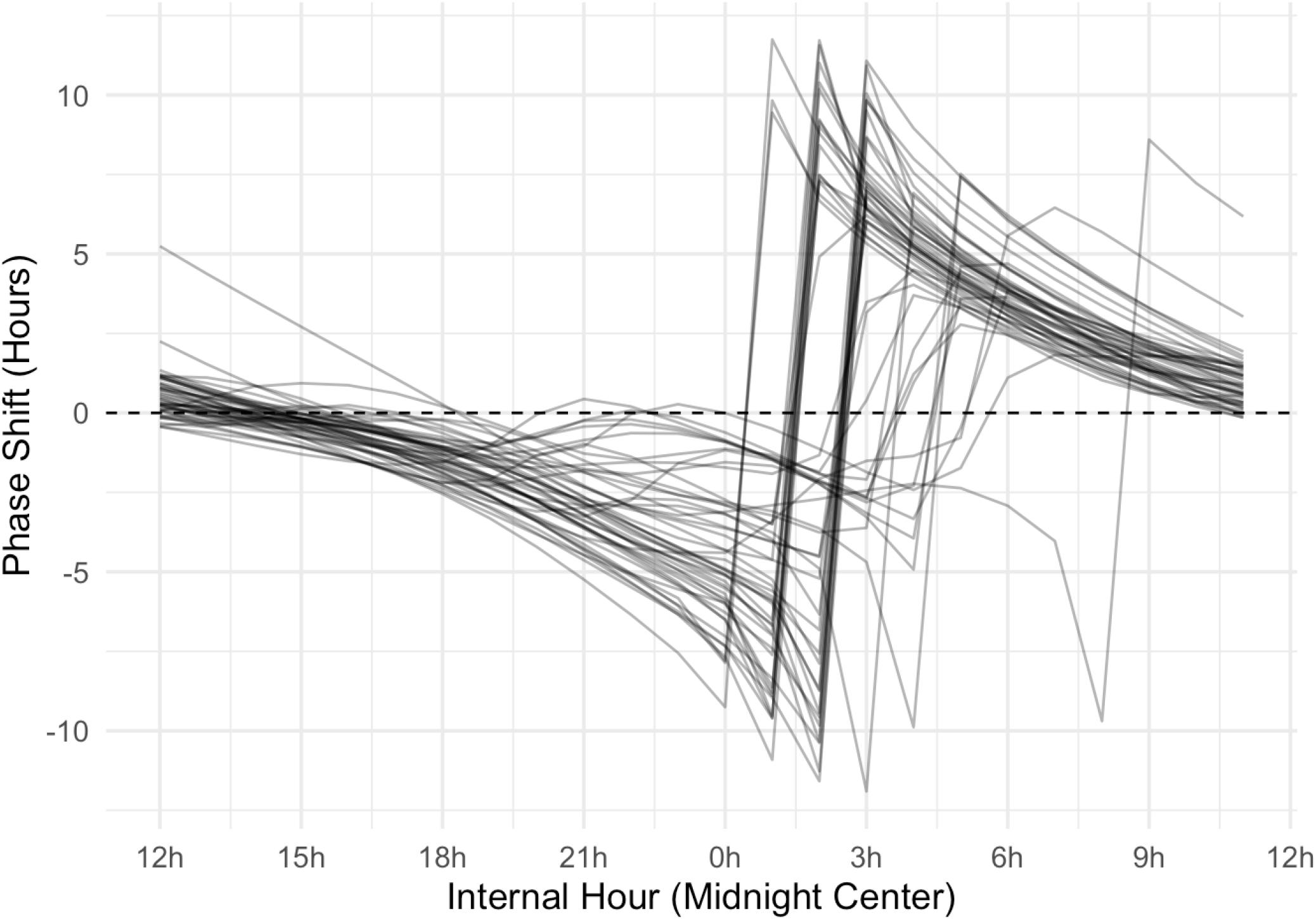
Individual phase response curves at 5,000 lux. Phase response curves generated for each participant (n = 59) using likelihoods estimated from their own recorded hourly light exposure, with the stimulus fixed at a one-hour pulse of 5,000 lux. The prior was the same for every participant (κ = 4). Each line represents one individual. The x-axis represents internal circadian time centred on midnight, and the y-axis shows the predicted phase shift in hours, where positive values indicate advances and negative values indicate delays. Despite the identical light stimulus, the timing of peak sensitivity and the amplitude of the shift vary across individuals, reflecting differences in the probability each person assigns to encountering 5,000 lux at different times of day based on their habitual light environment. Individuals whose typical light exposure includes brighter or later evening light show their peak sensitivity shifted later into the night relative to those with lower habitual evening light levels. Light exposure data were collected over one week in 59 UK adults using wrist-worn spectrometers^24^.

The same light stimulus can also have different phase-setting effects depending on light history, such as exposure during the day: high light levels during the day reduce the effect of light in the evening, and vice versa^35^. In our model, this can arise from the width of the prior. To illustrate this, we chose the participants with the lowest and highest mean illuminance between 07:00 and 18:00, the low and high daytime exposure cases (Figure 8). Each person’s light was summarised by this daytime mean and the mean log illuminance for the remaining hours. The model was run for two days on each person’s light, starting each day at 06:00. The clock’s phase was held at clock time, so that only the width of the prior changed. Between observations the prior slowly lost concentration, and at each hour it narrowed or broadened depending on how well the recorded light agreed with the light expected at that time of day. We then simulated a one-hour pulse of 31 lux at every hour of the day to give a PRC for each person (Figure 9). We used 31 lux because this is the light produced by an e-reader, and evening e-reader use has been shown to delay the circadian clock^36^, although whether this is biologically meaningful is debated. Both participants received the same pulse and the same likelihood, so they differed only in the width of the prior. For the participant with low daytime exposure, daytime light was far dimmer than expected, so it disagreed with the clock’s estimate, the prior broadened, and the pulse produced a large phase shift. For the participant with high daytime exposure, daytime light agreed with the clock’s estimate, the prior stayed narrow, and the same pulse produced almost no phase shift (Figures 9 and 10). These results illustrate a core prediction of the framework, that differences in circadian light sensitivity between people need not reflect differences in photoreceptor function or clock mechanisms, but may instead arise from differences in the learned statistical structure of each person’s light environment.

**Figure 8:**
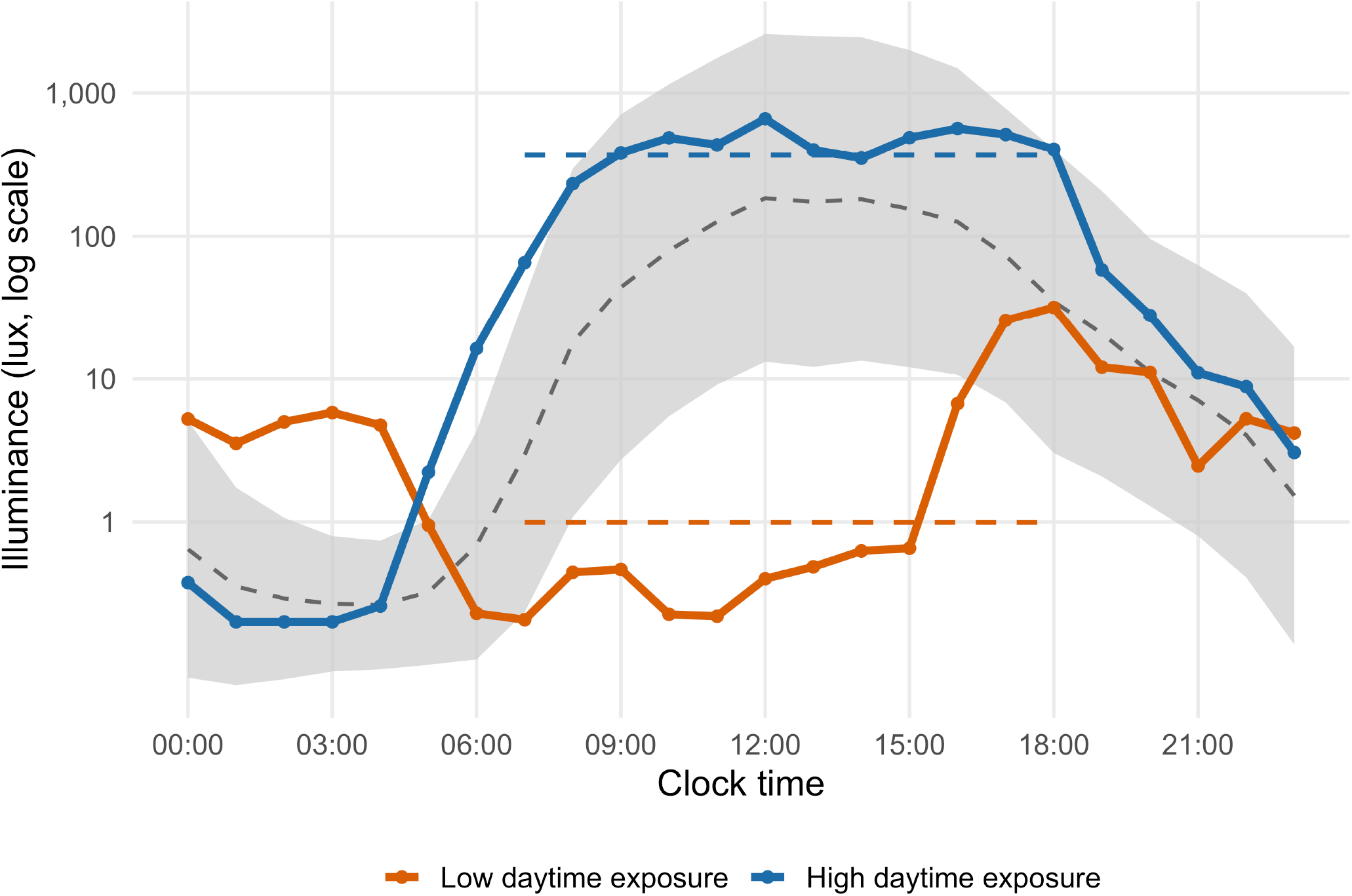
Daytime light of the two case participants. The x-axis is clock time and the y-axis is photopic illuminance (lux, log scale), shown at each hour of the day for the participant’s with the lowest (orange) and highest (blue) mean illuminance between 07:00 and 18:00 among the 59 UK adults recorded by Didikoglu et al.^24^. Each point is the participant’s mean log illuminance in that hour. The grey band is the mean ± 1 SD of log illuminance across all participant’s at that hour, and the dashed grey line is its mean. Together they give the light that the model expects at each time of day. The dashed coloured segments show each participant’s mean log illuminance between 07:00 and 18:00. The model uses this daytime mean, and the mean for the remaining hours, as the light of each participant across the day. The daytime light of the high exposure participant is close to or above the expected light, so it supports the clock’s estimate of the time and the prior stays narrow. The daytime light of the low exposure participant is far below the expected light, so it is improbable at the time the clock believes it to be, and the prior broadens. At each step the model multiplies the prior by the likelihood of the observed light, takes the circular mean and mean resultant length of the product, and returns the concentration κ with the same resultant length. Between steps κ decays slowly towards zero (time constant 240 h), which represents the clock’s uncertainty about its own tick.

**Figure 9:**
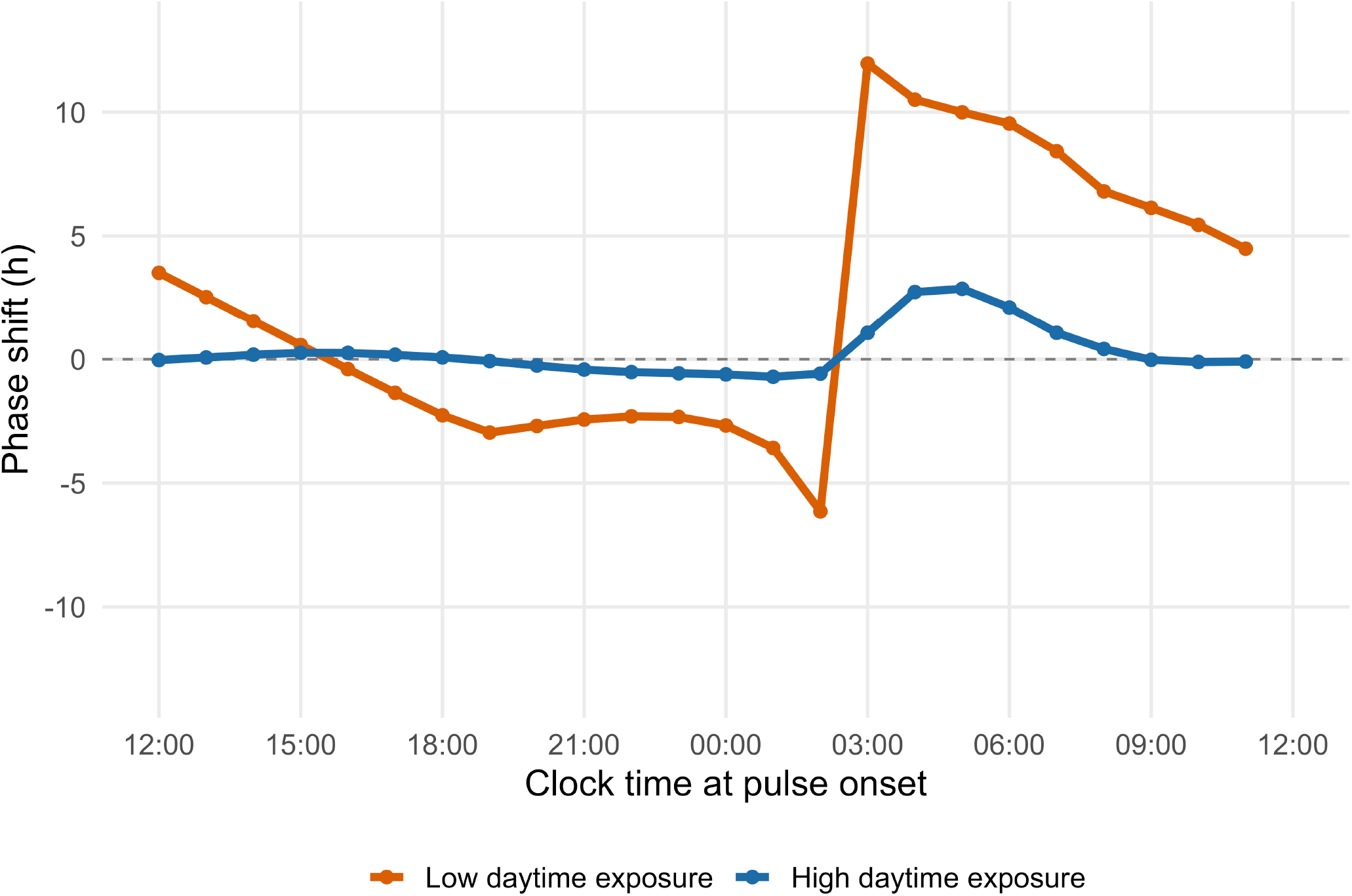
Phase response curves to a 31 lux pulse for the participants with low and high daytime exposure. Phase shift (hours) produced by the model for a one-hour pulse of 31 lux (the light of an e-reader) in the participant with low daytime exposure (orange) and the participant with high daytime exposure (blue) shown in Figure 8. The x-axis is the clock time at the start of the pulse, centred on midnight, and the y-axis is the phase shift in hours, with advances positive and delays negative. For each pulse onset, the model was first run for two days on that participant’s own light (Figure 8), which gave the concentration κ of the prior at that time. The clock was assumed to be at the participant’s clock time at the start of the pulse. The phase shift is the difference between the centre of the posterior after the one-hour pulse and the centre of the prior propagated forward over the same hour. The likelihood of 31 lux was estimated from the hourly distribution of illuminance in the 59 UK adults^24^, using a kernel density estimate of the log illuminance at each hour, and each hour of light counted as one hour of evidence. Both participant’s received the same pulse and the same likelihood, so they differ only in κ. The low exposure participant has a wider prior and shows a large phase shift. The high exposure participant has a narrower prior and shows almost no phase shift. The size of the shift at each time is approximately the slope of the log-likelihood of the pulse at that time divided by κ.

**Figure 10:**
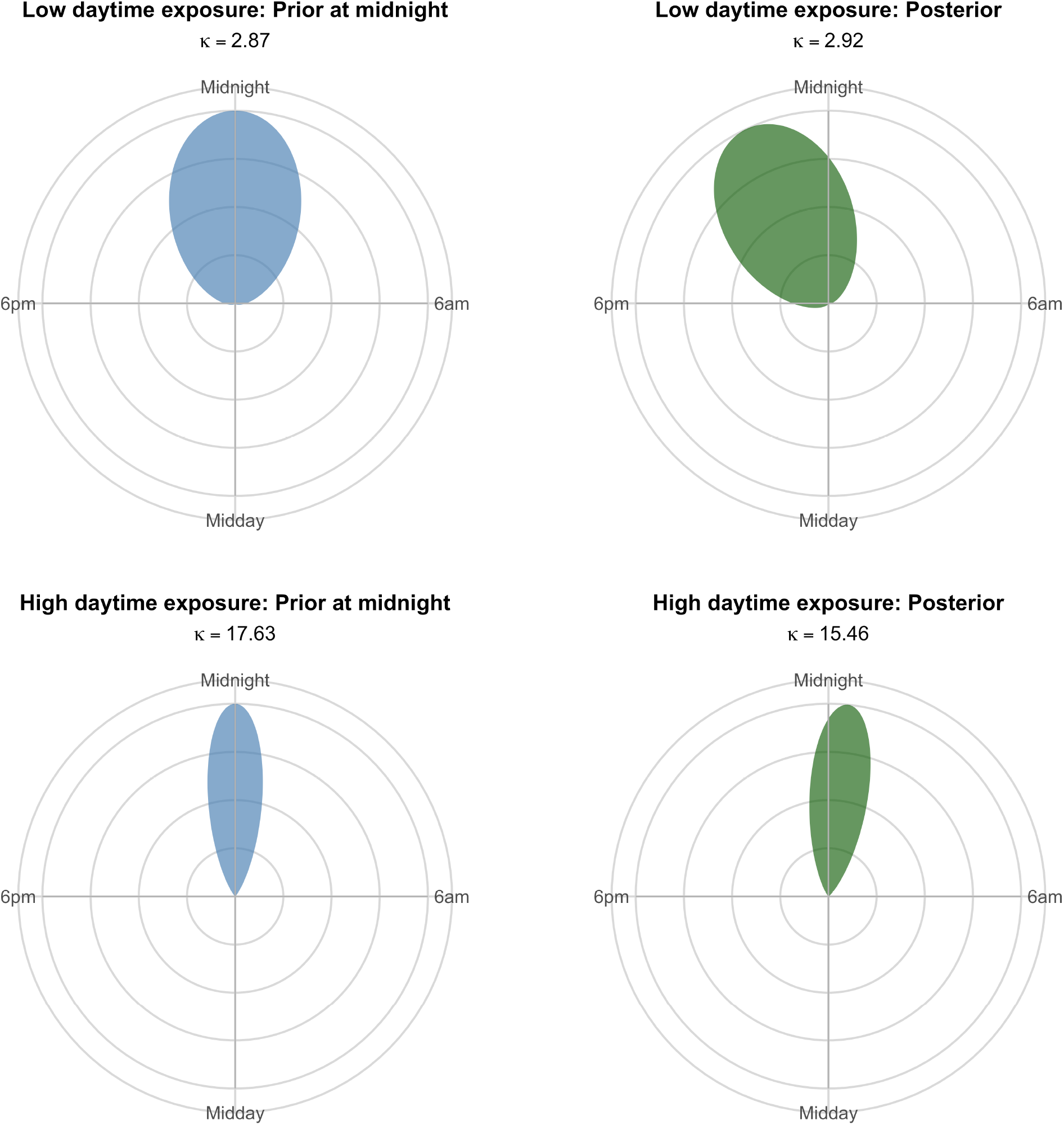
Prior and posterior of the clock for a midnight pulse in the participants with low and high daytime exposure. The clock’s belief about the time of day before (left, blue) and after (right, green) a one-hour, 31 lux pulse that begins at midnight, for the participant with low daytime exposure (top row) and high daytime exposure (bottom row). The angle of each panel is the clock’s estimate of the time of day (midnight at the top, 6am on the right, midday at the bottom and 6pm on the left). The height of each distribution is the probability the clock assigns to that time, scaled so that the peak of each panel reaches the same height. The prior (left) is a von Mises distribution centred on the participant’s clock time at midnight, with a concentration κ (printed in each panel) taken from the two-day simulation in Figure 8. The low exposure participant has a lower κ and so a wider prior, which is the clock’s reduced confidence after a day of light that disagrees with its expectation. The posterior (right) is the product of the prior and the likelihood of the 31 lux pulse over the hour, normalised and summarised as a von Mises distribution with the same circular mean and mean resultant length. At midnight 31 lux is improbable, and the likelihood is greater at earlier times in the evening, so the evidence tilts the estimate towards earlier times. The posterior of the high exposure participant stays close to its prior, because the narrow prior assigns little probability to the times that the evidence favours. The posterior of the low exposure participant moves further in the same direction, because its broad prior assigns more probability to those times. The difference between the centre of each posterior and the centre of its prior propagated forward over the same hour is the phase shift plotted at midnight in Figure 9.

## Combining The Features of Light: The Weighting of Light Features is Not Fixed

So far, we have modelled the central clock’s estimate of astronomical time using light intensity alone. Here we add spectral composition (the wavelength ratio) and the rate and direction of change of both, and show how the clock might use these features when they provide complementary or conflicting information. One of the core principles of the model is that the weight given to each feature of light is not fixed, but changes with the reliability of that cue at any given time. As we are asking what information is available in the light itself, the analyses in this section are performed using a flat prior, as is common in studies of weighted cue integration, and therefore focus on the likelihood. The likelihood is the probability of encountering an observed combination of light features given each candidate time of day. Because intensity and spectral composition naturally covary, their joint pattern often provides more information than either cue alone, allowing cue weighting to emerge from the statistical structure of the natural light environment.

We tested the model using the open dataset of Woelders et al.^37^, who continuously measured the spectrum of natural daylight over 100 days at a single location in Groningen, the Netherlands (53.24°N, 6.54°E). Because the position of the sun changes continuously throughout the day, and both the intensity and spectral composition of daylight vary systematically with it, solar angle provides a natural representation of astronomical time and therefore the quantity the circadian clock seeks to estimate from incoming light. The dataset comprised 88,960 spectral irradiance measurements (380–780 nm; 1 nm resolution) collected approximately every three minutes using a spherical (4π) diffuser that captured light arriving from all directions, closely approximating the natural light environment experienced by a stationary observer. Two photic cues were derived from each observation. Intensity was quantified as the log-transformed, lens-corrected melanopic photon flux, representing total light weighted by the spectral sensitivity of melanopsin (peak sensitivity approximately 480 nm). Spectral composition was quantified as the ratio of short-to long-wavelength cone excitation (S/L), derived from the MacLeod–Boynton chromaticity coordinates, capturing the characteristic blue–yellow spectral shifts accompanying changes in solar angle at twilight. Both measures were calculated using the CIE 170-1:2006 photoreceptor sensitivity functions. Solar angle and daily cloud-cover index (KNMI scale; 0–8 oktas, an okta being one eighth of the sky covered, so 0 is clear sky and 8 is complete overcast) were taken from the original dataset. The dataset was divided into 2° solar-angle bins and all subsequent analyses were performed at this resolution. For each solar-angle bin, we estimated the mean, standard deviation and covariance of intensity and spectral composition (Figure 11).

**Figure 11:**
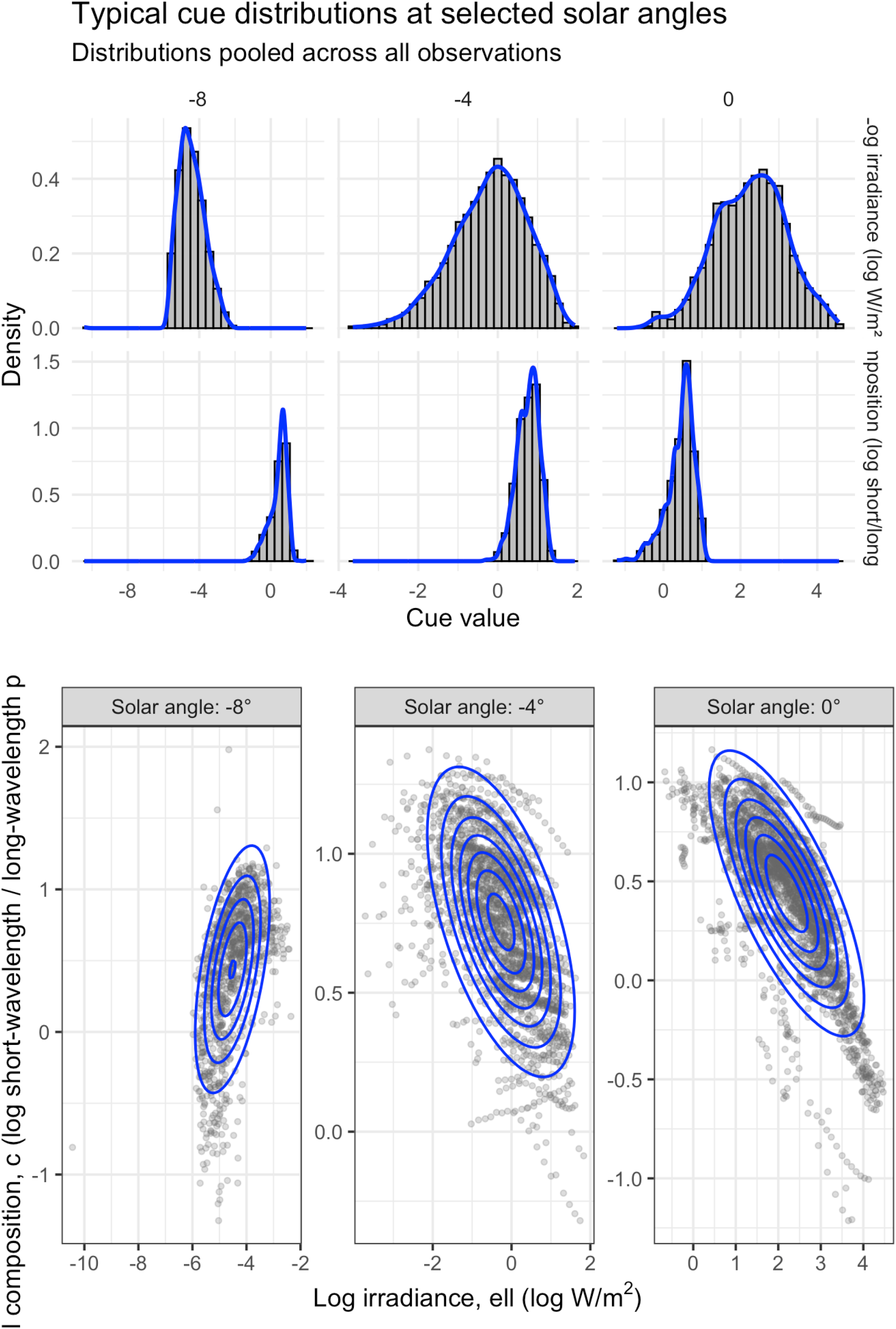
Joint distribution of intensity and wavelength at three solar angles. Scatterplots of intensity, the log-transformed melanopic photon flux, against spectral composition, the ratio of short-to long-wavelength cone excitation (S/L), at solar angles of −8°, −4° and 0°, derived from the Woelders et al.^37^ dataset. Each panel pools all observations within the 2° bin beginning at that solar angle (for example, −8° to −6°) across the 100-day measurement period. The x-axis is intensity (log melanopic photon flux) and the y-axis is spectral composition (S/L ratio). Blue contours show the fitted bivariate normal density at each angle. The location and shape of the joint distribution change systematically with solar angle. As the sun sinks further below the horizon, intensity falls and the light becomes relatively richer in short wavelengths, reflecting the blue colour of twilight. The covariance between the two cues, captured in the off-diagonal terms of Σ(*s*_*t*_) in the observation model, also changes with solar angle, so the joint pattern of intensity and wavelength provides evidence about astronomical time beyond that carried by either cue alone.

### Combining cues increases the stability of solar angle estimation

If the central clock integrates multiple features of light to arrive at a more reliable estimate of astronomical time, our model offers a formal description of how it might weight these features using the information available in natural light. To show this, we used our Bayesian clock model with a flat prior, so that its estimate was determined by the likelihood alone. We first built a reference distribution for every 2° solar-angle bin between −6° and 60°. To do this, we averaged the light recorded within each bin on each day, and then calculated the mean and variance of these daily averages across the dataset. Each reference distribution therefore describes the typical light at that solar angle and how much it varies from day to day. To test whether combining intensity and wavelength improves the prediction of solar angle compared to intensity alone, we then took 200 test observations at each true solar angle. Each test observation was the average light recorded within that bin on a single day, with the day chosen at random from the dataset. Each observation was compared with every reference distribution, once using intensity alone and once using intensity and wavelength combined, calculating how compatible the observation was with each, i.e. its likelihood. These likelihoods were normalised to give a posterior probability, and the posterior-weighted average was taken as the models estimate of the solar angle at which the observation was recorded. When an observation closely matches only one or two neighbouring reference distributions, the weighting is concentrated there and the estimate lands close to the true value. When it is similarly probable across many neighbouring distributions, the weighting is spread more broadly, and the resulting estimate is less accurate and less consistent from one observation to the next. Because the true solar angle of every observation is known, we calculated the error between each estimate and the true angle, and measured how tightly the 200 errors at each angle clustered using the interquartile range (IQR). A smaller IQR indicates that the model gave a similar answer across repeated observations from the same true solar angle, whereas a larger IQR indicates that its answers varied more from one observation to the next. Across the whole tested range, the errors of the intensity-only estimates were, on average, 1.65 times more spread out than those of the combined estimates (Figure 12). Combining intensity and wavelength improved consistency over intensity alone from around 0° up to around 28° solar angle, with the improvement largest at 14°, where the errors of the intensity-only estimates were 5.46 times more spread out than those of the combined estimates.

**Figure 12:**
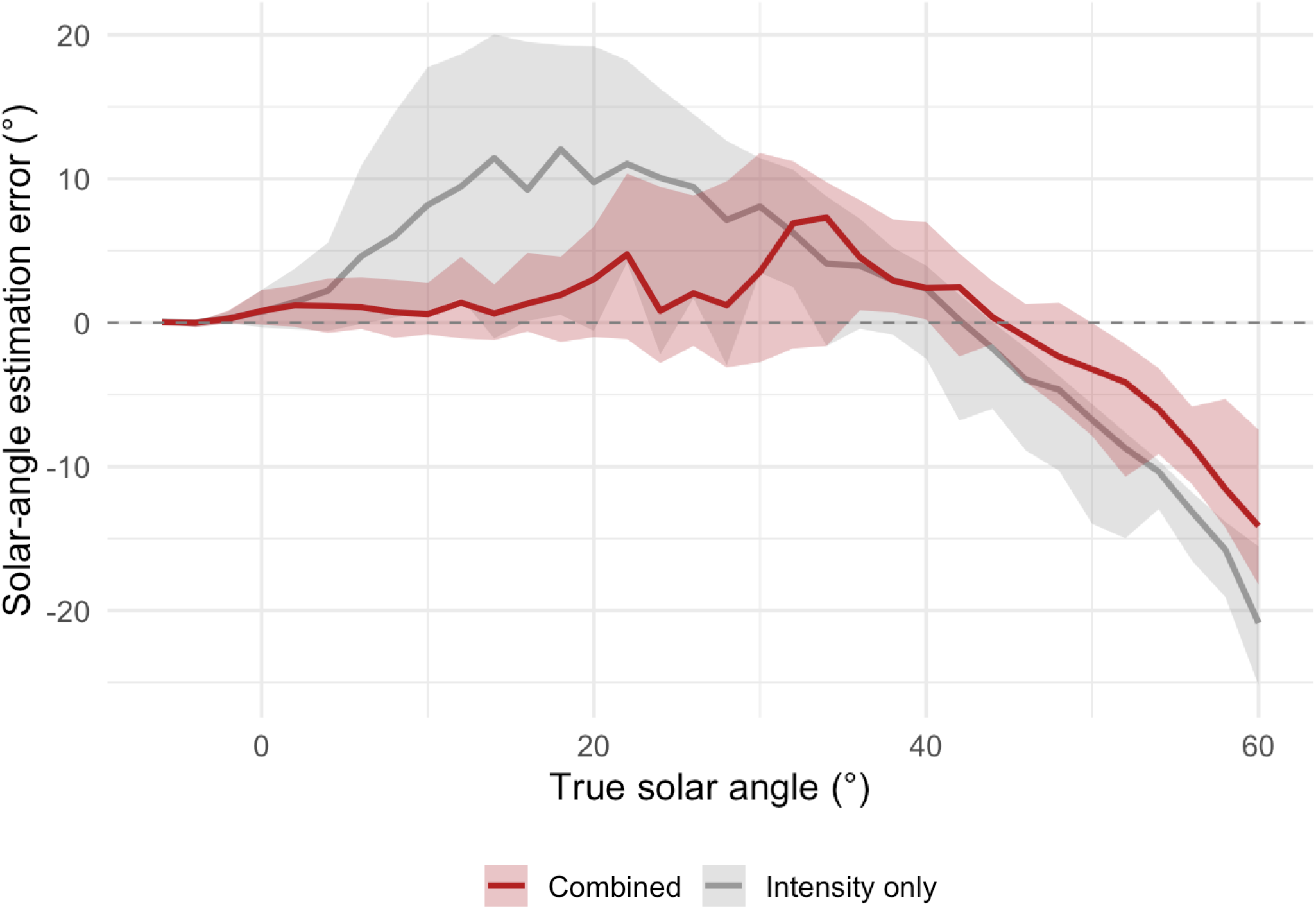
Stability of solar-angle inference, intensity alone versus combined cues. Median solar-angle estimation error (line) and interquartile range, IQR (shaded band), across repeated observations at each true solar angle, for intensity alone (grey) and the combined intensity-wavelength model (red). The x-axis is the true solar angle (°) and the y-axis is the error between the models estimate and the true solar angle (°). The IQR band shows the range spanning the middle 50% of the estimation errors at each true angle: a narrower band indicates that repeated estimates cluster more tightly around the same value, reflecting more consistent selection of the correct solar angle under naturally varying light conditions. The intensity-alone band is visibly wider than the combined band across most of the range, consistent with a mean IQR 1.65 times larger for intensity alone than for the combined model, with the largest difference at 14° solar angle (ratio 5.46). The dashed horizontal line marks zero error.

Alongside consistency, we measured a second property of the model, which we call gain. Gain captures how much the model’s estimate of solar angle changes when the input light changes by a fixed amount. We defined gain as the change in the model’s inferred solar angle produced by a one-standard-deviation increase in intensity, relative to the typical intensity observed at that solar angle, with the estimate based either on intensity alone or on intensity and wavelength combined. Gain is high when the light expected at neighbouring solar angles differs noticeably, so that even a small change in the observed light produces a large change in the models estimate, since the model can determine more accurately which angle the new light belongs to. Gain is low when the light expected at neighbouring solar angles is similar, so that the same change does little to shift the models estimate, since the model cannot use the light to tell the angles apart. Adding wavelength can raise gain, because neighbouring angles with similar intensity may still differ in wavelength. Within the informative twilight range, −6° to 12° solar angle, mean gain was 5.98 using intensity alone and 6.36 using intensity and wavelength combined. Combining intensity and wavelength therefore increased the model’s responsiveness to real changes in the light environment, in addition to the improvement in consistency reported above. This illustrates how the central clock might use the information in the light environment: by combining intensity and wavelength, it may obtain a more consistent and more responsive estimate of astronomical time during twilight.

Interestingly, beyond around 28°, combining wavelength with intensity no longer improved consistency, and in several bins the errors of the combined estimates were more spread out than those of the intensity-only estimates. The data therefore revealed a statistical dead zone, similar to the dead zone found in phase response curve experiments, the period during the subjective day in which light produces little or no phase shift^30^. Our analysis showed that mean intensity increased from 16.35 to 19.53 log photon units between −6° and 10° solar angle, but changed by less than 1 log unit (19.63 to 20.30) from 12° to 60°. Mean spectral composition decreased from 0.0658 to 0.0384 between −6° and 10°, but then remained within a narrow range (0.0380 to 0.0389) from 12° to 60°. At the same time, the covariance between intensity and spectral composition fell from a maximum of −0.0034 near 4° to near zero (−0.0000832) by 60°. Thus, beyond 12° solar angle, changes in the mean intensity and spectral composition between neighbouring angles became small relative to the standard deviation at a single angle, causing the natural variability at one angle to overlap with the typical values at neighbouring angles. The model therefore began to lose discrimination between neighbouring solar angles, with the benefit of combining wavelength and intensity disappearing by around 28°. At higher solar angles, the cues could no longer reliably distinguish one angle from another, so light in the middle of the day carried little information about time of day. The highest solar angles (40°–60°) occurred only during the summer months at this latitude and were represented by a smaller sample of 36 days.

### Reliability-weighted integration under cloud cover

If the central clock weights each feature of light according to how reliably it signals astronomical time, the weight given to wavelength should change when conditions alter the reliability of intensity. Cloud cover provides a natural test of this. Clouds scatter and absorb sunlight, so the intensity of light at a given solar angle varies more from day to day under cloudy skies, whereas the spectral composition of twilight is thought to be less affected^37^. Consistent with this, wavelength signals support circadian entrainment in mice when light intensity is a weak or ambiguous indicator of time of day^16^. We therefore asked whether the weight our model assigns to wavelength increases with cloud cover. The analysis was restricted to twilight (−6° to 12° solar angle). Daily cloud cover (0–8 oktas) was treated as a continuous variable. For each cloud level from 1 to 8 oktas, we built a reference distribution for every 2° solar-angle bin from all days, giving each day more weight the closer its cloud cover was to that level. For each cue, we then calculated its reliability: how much its mean changes between neighbouring solar angles relative to its day-to-day variability, expressed as uncertainty about solar angle in degrees.

This analysis follows the design of classic cue-integration experiments used to test Bayesian models of perception. In these experiments, noise is added to one cue to reduce its reliability, and a small conflict between cues is introduced to measure how much weight each receives; reliability-weighted integration predicts that the weight shifts away from the noisier cue^27^. Here, cloud cover provides the noise naturally, by making intensity more variable from day to day, and cue conflict is used to measure the resulting weights. The conflict is necessary because, in natural light, intensity and wavelength almost always indicate the same solar angle, so the model’s estimate lands in the same place whichever cue it relies on most, and the contribution of each cue is hidden. Pairing cues that indicate different solar angles reveals this contribution: where the estimate lands between the two indicates how much weight each cue received. We therefore paired an intensity observation from a day at the test solar angle with a wavelength observation from a day 6° or 10° lower. The wavelength weight was calculated as how far the model’s estimate moved from the intensity-only estimate toward the wavelength’s solar angle, as a proportion of the distance between them. A weight of 0 means intensity alone determined the estimate, and a weight of 1 means wavelength determined it. Because cloud cover varies with season, each trend was tested against a null distribution in which cloud cover was shuffled between days within the same month (200 permutations).

Clouds reduced the reliability of intensity. Uncertainty about solar angle from intensity doubled from 0.95° at 1 okta to 1.96° at 8 oktas (p = 0.005), whereas uncertainty from wavelength showed no consistent trend with cloud cover (range 3.65° to 7.31°; p = 0.21). Intensity therefore remained the more reliable cue, but its advantage over wavelength narrowed as cloud cover increased. The model responded by giving wavelength more weight (Figure 13). With wavelength taken from 6° lower, the wavelength weight rose from −0.19 at 1 okta, where few clear days were available, to 0.44 at 8 oktas (p = 0.005); with wavelength taken from 10° lower, it rose from 0.07 to 0.52 (p = 0.04). As expected for a Bayesian observer, even when the cues disagreed, combining them reduced uncertainty about solar angle in 87.2% of trials for the 6° conflict and 97.4% for the 10° conflict, with mean reductions in entropy of 0.44 and 0.84 nats. Entropy here is the Shannon entropy of the posterior, which measures how broadly probability is spread across candidate solar angles, in nats, the unit of information given by the natural logarithm. A reduction of about 0.69 nats corresponds to halving the number of solar angles that remain plausible. Together, these results offer an explanation for why wavelength becomes more useful under cloud: clouds degrade the reliability of intensity, and a clock that weights cues by their reliability shifts its weight toward wavelength accordingly. The relative contribution of each feature of light is therefore set by the prevailing conditions.

**Figure 13:**
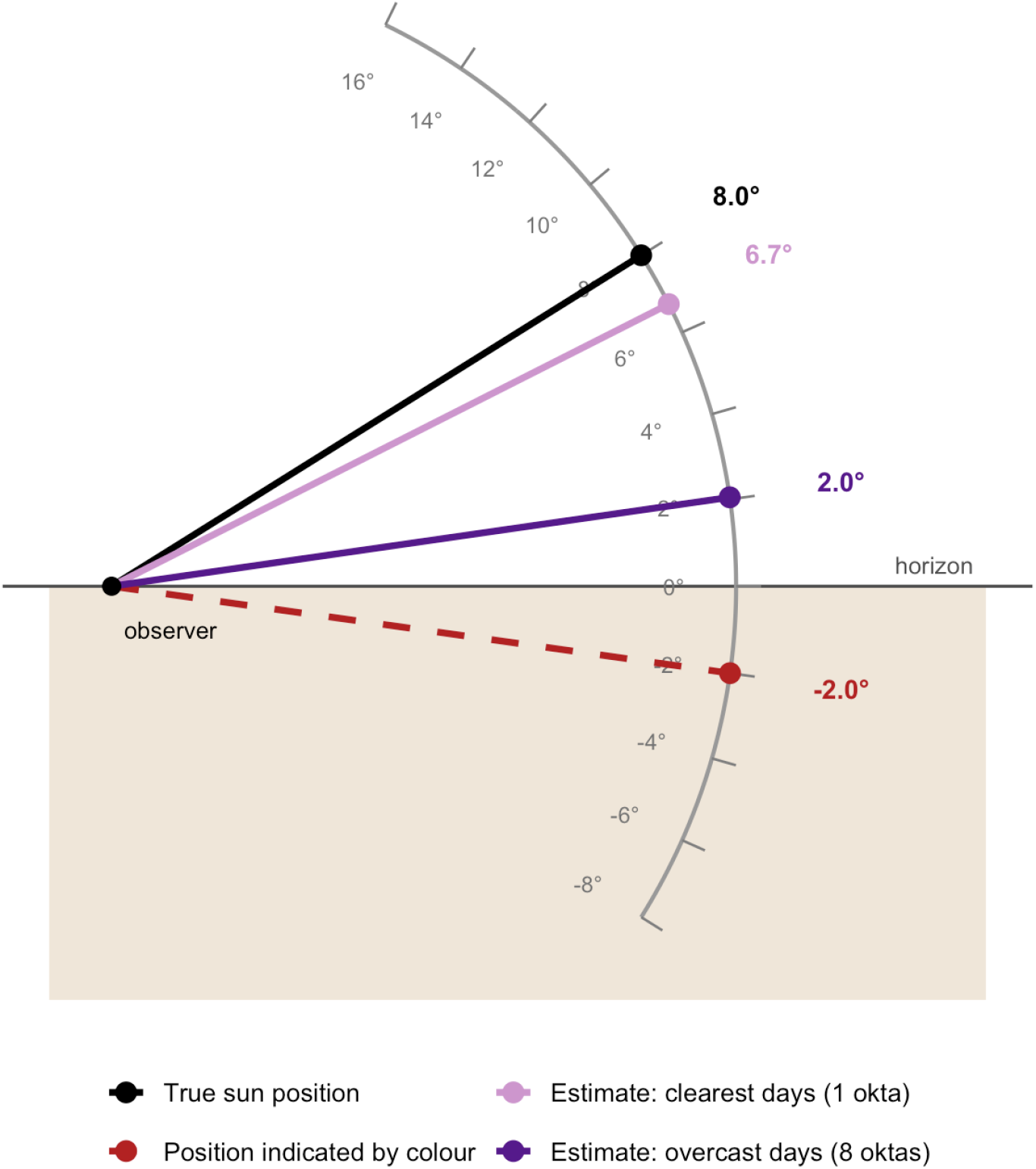
Cloud cover shifts the model’s estimate of the sun’s position toward wavelength. The diagram shows a view of the sky from an observer (black point) standing on the ground (shaded), looking toward the horizon (horizontal line). The grey arc represents the sky, marked with solar elevation in degrees; positions above the horizon line are positive solar angles, and positions below it are negative solar angles, when the sun is below the horizon. Each line runs from the observer to a position on the arc, representing an estimated or actual position of the sun. To test how the model weights intensity and wavelength, the two cues were placed in conflict. An intensity observation recorded at a true solar angle of 8° was paired with a wavelength observation recorded at −2°, 10° lower, so that the two cues indicated different positions of the sun. In natural light, intensity and wavelength almost always indicate the same solar angle, which hides how much the model relies on each; separating them in this way reveals the weight each cue receives. The black line shows the true position of the sun, indicated by intensity (8°). The red dashed line shows the position indicated by the conflicting wavelength (−2°). The purple lines show the model’s estimate of solar angle when it receives the two conflicting cues together, on the clearest days (1 okta; light purple) and the most overcast days (8 oktas; dark purple), with each estimate labelled on the arc. The closer an estimate lies to the black line, the more the model relied on intensity; the closer it lies to the red line, the more it relied on wavelength. On the clearest days, the estimate lies close to the true sun position, because intensity is a reliable indicator of solar angle under clear skies and the model weights it heavily. On overcast days, the estimate moves toward the position indicated by wavelength. Clouds make intensity more variable from day to day, reducing its reliability, while the reliability of wavelength shows no consistent change, so the model shifts its weight toward wavelength. The figure illustrates a single example solar angle; the change in wavelength weight with cloud cover across all twilight angles is reported in the text.

### Resolving the dawn–dusk ambiguity

Instantaneous measurements of light intensity and spectral composition are intrinsically ambiguous because similar values occur twice each day, once during dawn and again during dusk. They indicate the elevation of the sun, leaving open whether it is rising or setting. Within our model, this ambiguity is resolved by the direction of change of light intensity. Throughout this section, rate of change refers to the rate of change of intensity. During the morning, light intensity increases and the spectrum shifts away from the blue-enriched light of twilight as the sun rises. During the evening, intensity decreases and the spectrum becomes progressively bluer as the sun sets. The direction of change therefore identifies whether the observer is in the morning or evening half of the solar day. To show how the clock might use this information, we used the model to decide whether each observation came from the morning or the evening. For each reading, we calculated the rate of change of intensity as the difference between that reading and the reading taken 15 minutes earlier, divided by the time between them. The sign of this value indicates whether light is increasing or decreasing, and its size indicates how quickly. For every 2° solar-angle bin between −14° and 14°, a range reached on every day of the dataset, we then built separate reference distributions for morning and evening from the daily averages of intensity, spectral composition and the rate of change of intensity. The rate of change of spectral composition was not included, as the direction of change in intensity alone resolved the ambiguity. Each observation was compared with the morning and evening reference distributions at its solar angle and assigned to whichever gave the higher likelihood. So that each observation was always compared with reference distributions built from other days, each day was left out in turn, the reference distributions were built from the remaining days, and the left-out day was then classified. This was done first using intensity and spectral composition alone, and then adding the rate of change of intensity. Using intensity and spectral composition alone, classification accuracy was 54.1%, indistinguishable from chance, confirming that these features alone cannot distinguish dawn from dusk. Adding the rate of change increased overall classification accuracy to 89.9%, with perfect performance (100%) between −8° and 0° solar angle. Accuracy declined towards the edges of this range, falling to 65–79.5% at around −14° and at 12–14°, where intensity and spectral composition became progressively less distinctive. Calculating the rate of change over 15 minutes, rather than between consecutive readings 3 minutes apart, increased classification accuracy modestly (89.9% versus 87.8%), reflecting the reduced influence of short-term fluctuations in natural light. At high solar angles (40°– 60°), which were reached only on summer days, accuracy with the rate of change was 53.4%, close to chance, consistent with the statistical dead zone described above.

To illustrate how this works for a single observation, we selected a real observation recorded at dusk on 23 February 2016 at 16:56, when the sun was at −0.4° solar angle. It was chosen from evening observations between −4° and 0° as one with the steepest fall in intensity, providing a clear example of the direction-of-change cue. At this solar angle, intensity and spectral composition are nearly identical to their values during the corresponding morning twilight, so on the basis of these features alone the observation could plausibly be mistaken for dawn. Using reference distributions built for each hour of the day, we calculated the probability of this observation at every hour as each feature was added to the model in turn: intensity alone, then intensity with wavelength, then the size of the rate of change, and finally its direction (Figure 14). With intensity alone, probability was split almost equally between dawn and dusk. Adding wavelength and the size of the rate of change narrowed the estimate around both twilights, and only the direction of change removed the dawn peak, identifying dusk as the correct time. Different properties of natural light therefore inform different aspects of astronomical time. Intensity and spectral composition indicate the elevation of the sun, and the direction of change identifies whether it is rising or setting. Together, these features would allow the clock to estimate astronomical time unambiguously.

**Figure 14:**
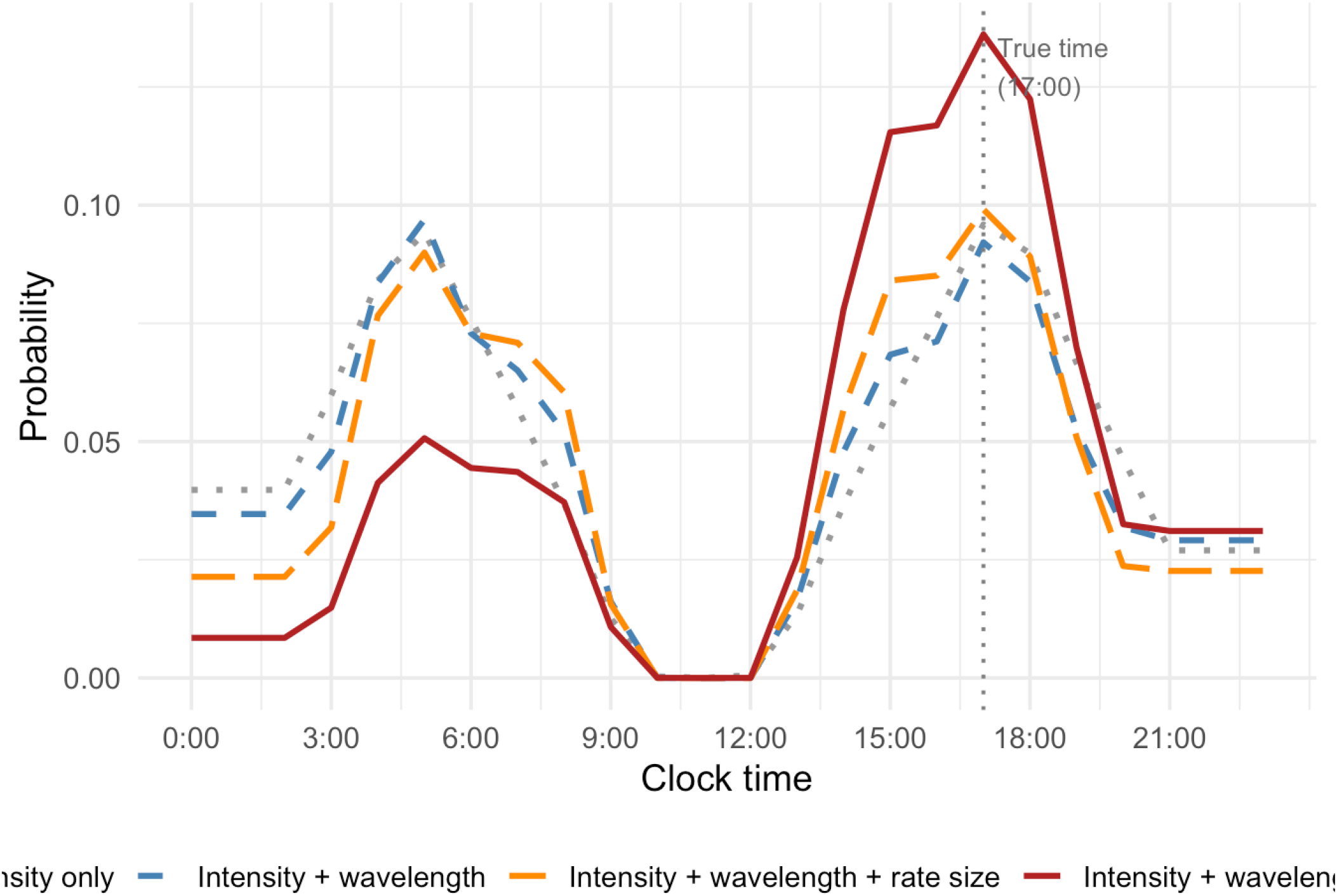
Progressive constraint of time-of-day estimation by multiple light features. Probability of a real dusk observation (23 February 2016, 16:56, solar angle −0.4°; data from Woelders et al.^37^) across the 24-hour day under four successive combinations of cues. The x-axis is clock time and the y-axis is the probability of the observation at each hour, normalised to sum to one. With intensity alone (grey dotted), probability is nearly equally split between dawn and dusk, reflecting the fundamental ambiguity that the same melanopic intensity occurs at both twilight transitions. Adding wavelength (blue dashed) and the magnitude of the rate of change (orange, long dashed) progressively narrows the distribution but does not resolve the ambiguity — both dawn and dusk peaks increase together. Only when the direction of change is included (red solid) — whether intensity is rising or falling — does the ambiguity resolve, with the dawn peak collapsing and the dusk peak identified as the correct branch. The dotted vertical line marks the true clock time of the observation (17:00). Shannon entropy decreases progressively across the four conditions (2.41, 2.29, 2.16, 2.06 nats), reflecting the reduction in uncertainty as each cue is added. For illustration, the standard deviation of the rate-of-change cue was inflated threefold and each curve was smoothed with a five-hour moving average.

Taken together, these analyses show that each feature of natural light contributes to the clock’s estimate of astronomical time in a way that depends on the conditions under which it is observed. Combining intensity and wavelength improves the estimate during twilight but adds little in the middle of the day; the weight given to wavelength increases when cloud cover reduces the reliability of intensity; and the direction of change resolves the ambiguity between dawn and dusk that intensity and wavelength alone cannot. The weighting of the features of light is therefore set by the solar angle, the weather and the time of day at which light is observed, rather than fixed in advance.

### Reliability-weighted Integration under cue conflict

Here we use the same cue-conflict design across all well-sampled solar angles, regardless of cloud cover, pairing an intensity reading with a spectral composition reading from a day 6° or 10° lower. For each conflicting observation, the weight given to spectral composition was calculated as how far the model’s estimate moved away from the estimate produced with the correctly paired, non-conflicting spectral composition, toward the conflicting cue’s angle, as a proportion of the distance between the two cues’ angles. We also measured the Shannon entropy of the posterior, which indicates how precisely, rather than how accurately, the model estimates astronomical time.

Wavelength weight and entropy both became unstable beyond approximately 30° solar angle, because these high elevations occur only during the summer months at this latitude and are represented by a much smaller number of observations. We therefore restricted this analysis to the well-sampled range between −6° and 28° solar angle. With a 10° cue conflict, the model’s estimate deviated by a mean of 6.19° from the non-conflicting estimate, and spectral composition contributed a mean weight of 0.407 to the combined estimate. With a 6° cue conflict, the mean deviation was 5.44°, and spectral composition contributed a mean weight of 0.291. At the 10° conflict, entropy decreased by a mean of 0.606 nats; at the 6° conflict, entropy decreased by a mean of 0.336 nats. This decrease occurred in the majority of observations, 80.2% at the 10° conflict and 71.1% at the 6° conflict, indicating that, as expected for a Bayesian observer, combining conflicting cues produced a more precise estimate than intensity alone, even when the two cues disagreed.

The weight assigned to spectral composition varied systematically across the day. It was negligible near −4° solar angle, increased rapidly through early twilight, and reached a plateau between approximately 8° and 20° solar angle, where the mean weight remained between 0.68 and 0.80. Beyond this plateau, the weight assigned to spectral composition declined, falling to 0.485 at 26° and 0.469 at 28°. Spectral composition received the most weight when the conflicting wavelength came from twilight, where it is most informative, and intensity at the true angle was becoming less so. Together, these results show that the clock does not treat intensity and spectral composition as fixed, equally weighted inputs, but instead combines them according to how reliably each one signals astronomical time at that particular point in the day.

## The Duration of a Light Pulse and the Saturation of a Phase Shift

The duration of a light stimulus affects the phase response of the central clock, with longer pulses shifting the clock more than shorter ones^38^, but not in proportion to their length. Per minute of exposure, a short pulse of bright light can be several times more effective than a long one^39^. This levelling off is called the saturation effect. In our model, it emerges from the accumulation of evidence over time. As evidence accumulates, the clock’s estimate moves towards the time that the light implies, the prediction error is reduced and the prior narrows, so each further update produces a smaller shift and the total shift levels out.

To show how the phase shift saturates with the duration of a light pulse, and how this interacts with the intensity of light, we gave the model clock a constant light pulse of 1, 10, 100, 1,000 or 10,000 lux for 3 hours, starting at 03:00 (Figure 15). The times of day at which each light level is most likely came from the hourly light recordings of 59 UK adults^24^. The clock began with the same prior for every light level. At every moment the clock combined its prior with the light it saw, and the posterior became the prior for the next moment, so the evidence accumulated over the pulse. Light close to what the clock expects at 03:00, such as 1 lux, hardly moved the clock. Light of 10 lux, which is most likely in the evening, moved the clock slowly at first and was still shifting it when the pulse ended. Brighter light is typical of the day and unlikely at 03:00, so its likelihood points far from the clock’s current estimate. This gave a large prediction error, a fast shift and then a levelling off, with brighter light shifting the clock faster. The saturation effect therefore emerges directly from the recursive update, without any additional rule or mechanism, and may reflect an adaptive strategy that prevents excessive or unstable adjustments to continuous or repeated light. The same structure also protects the clock against transient, unreliable signals, such as a sudden increase in intensity when a cloud passes and direct sunlight breaks through. A brief event contributes less evidence than sustained light, so large phase shifts require consistent evidence that repeatedly reinforces the same prediction error.

**Figure 15:**
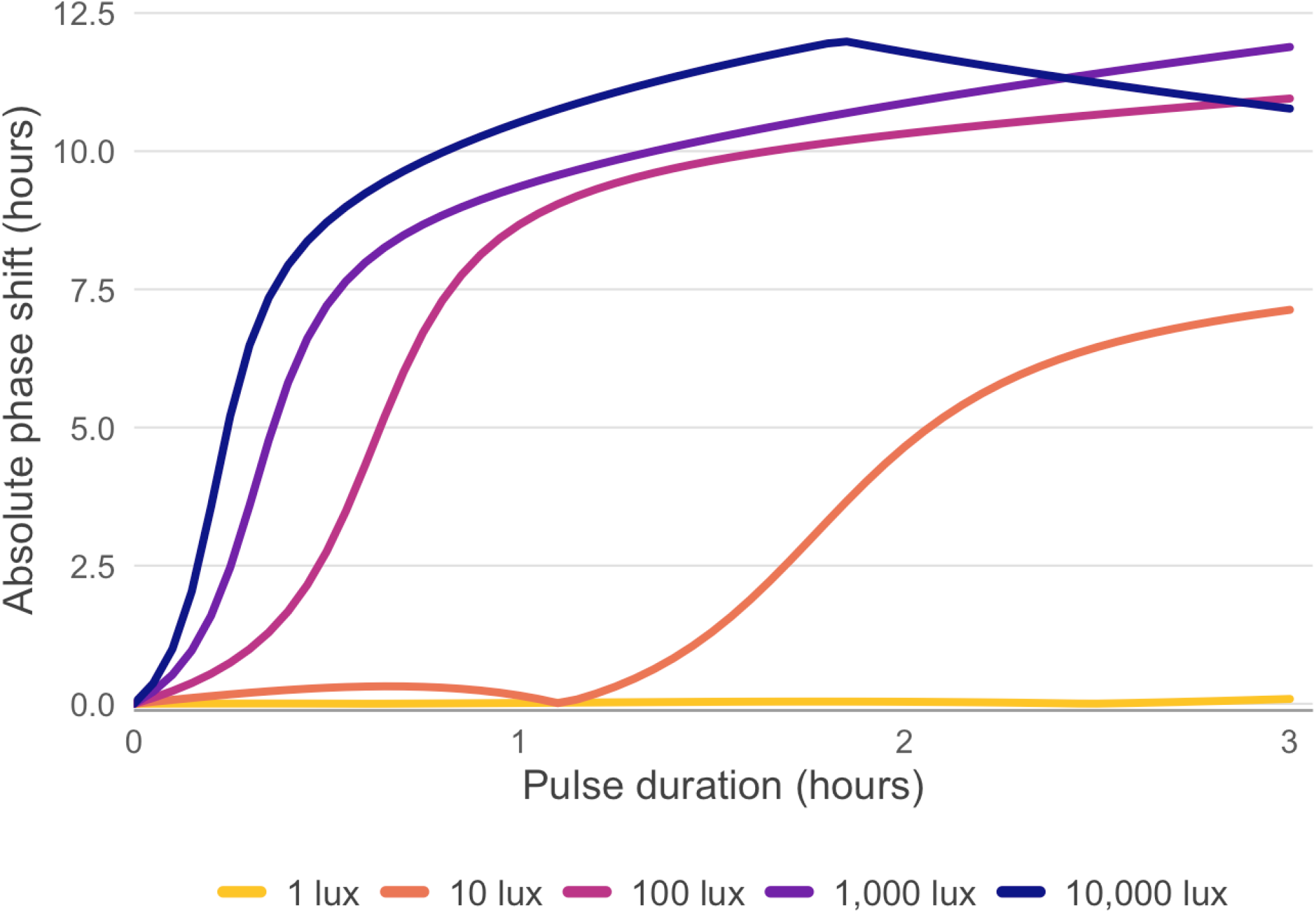
Phase shift saturates with the duration of a light pulse, and the speed and size of the shift depend on light intensity. Absolute phase shift (hours) produced by the Bayesian observer model for constant light pulses of 1, 10, 100, 1,000 and 10,000 lux (colour, from yellow for the dimmest to dark purple for the brightest) as a function of pulse duration. The x-axis is the duration of the light pulse in hours, from 0 to 3 hours. The y-axis is the absolute phase shift in hours, which is the difference from an identical clock that received no light, whether the shift is an advance or a delay. The largest possible value is 12 hours. Every pulse began at 03:00. The times of day at which each light level is most likely came from the hourly photopic illuminance recorded in 59 UK adults^24^. As evidence accumulates, the clock’s estimate moves towards the time that the light implies. The clock and the observation become increasingly aligned, the prediction error is reduced and less phase shift is needed, so the total shift levels out. Light of 1 lux points to a time near the clock’s estimate, so it moves the clock hardly at all. Light of 10 lux moves the clock slowly at first and was still rising at 3 hours. Light of 100 lux and brighter is typical of the day and unlikely at 03:00, so it moves the clock most of the way within the first hour and then levels off.

## Discussion

Circadian photoentrainment may be understood as Bayesian inference, in which the central clock, the suprachiasmatic nucleus, maintains a probabilistic estimate of astronomical time that is updated using sensory evidence from the natural light environment. Phase shifts emerge from the mismatch between the clock’s internal estimate of expected light, given its internal phase, and the actual observed values of light, a prediction error, while the magnitude of those shifts depends on the uncertainty associated with both, combined as a weighted average, with the updated phase closer to the estimate that is most reliable. Previous mathematical models of central clock function have successfully described circadian oscillators and phase resetting to light^19–22^. These models generate predictions that, in many cases, are aligned with ours and, under controlled laboratory conditions, would produce similar results. However, we suggest that our framework contributes an additional principle: the clock’s response to light depends on prior light history, and therefore does not respond to the absolute value of a light stimulus but is modulated by the typical range of light values experienced given the time of day. Indeed, perception is relative and based on comparisons, rather than on the encoding of absolute physical values^33,40^. However, rather than these adaptive processes unfolding across minutes and hours, as in other perceptual systems, the circadian response to light can be influenced by prior light exposure across many days. Based on Bayesian principles, and using data collected by 59 UK adults wearing a wrist-worn light sensor, we showed how the canonical shape of the phase response curve^30^ can emerge from the probability of encountering light at different phases of the circadian cycle. Because the probability of encountering a bright light, in our example 5000 lux, is low in the middle of the night, light of this magnitude generates a large prediction error and therefore large phase shifts. The dead zone follows the same logic in reverse. In our model, bright light during the subjective day carries little information because it does not vary in a way that distinguishes one time from another. Conversely, under our model, a dark pulse during the middle of the subjective day is also informative, indicating the clock’s internal estimate of astronomical time may be inaccurate. Our model showed how a light of 1 lux produced opposite phase adjustment behaviour to brighter light, indicating that the dead zone may not be a period of absolute insensitivity, but insensitivity to expected light. Indeed, phase shifts to a dark pulse have been observed in both rodents and humans^41–43^. Phase response curves, however, are measured in animals free-running in constant darkness^30^, where no light is being sampled and yet the dead zone is still expressed, so natural light statistics cannot be what generates it online. They may instead explain why it exists: because light in the middle of the subjective day is a poor discriminator of time, natural selection would favour a clock that discounts it, internalising those statistics as an innate prior rather than one acquired within a lifetime. Our framework accommodates both, with ancestral light statistics setting the shape of the response and prior light history modulating sensitivity within an individual. Using this logic, the model explains why the same light intensity can produce different phase responses across individuals. What is bright to one person at 8pm may be considered dim to another, assuming they have different light histories, which we showed by fitting PRCs to individual light exposure data^24^. This can help explain the shape of the human phase response curve to light, which remains contested. Whether humans possess a real dead zone, whether dim light from e-readers has a meaningful effect in the evening, and the maximum magnitude of phase advances and delays remain open questions. In our model, these contested findings are not mutually exclusive but reflect different combinations of the clock’s internal state, prior light history, and variability in the natural light environment. One of the core features of our model is that the integration of light cues depends on their reliability. Using light data collected by a stationary light sensor, recording natural light for 100 days in the Netherlands, we showed that light information regarding solar angle becomes more informative when these components are combined in a reliability-weighted average. Our model showed how the wavelength ratio of light may provide more reliable evidence about astronomical time under conditions of heavy cloud cover.

The integration of the intensity and wavelength of light leads to an unresolved anatomical question. Changes in spectral composition at twilight help stabilise photoentrainment, and single-unit recordings from the mouse SCN reveal two functionally distinct neuronal populations: those driven primarily by melanopsin-mediated irradiance signals centred around 480 nm, and those integrating short- and long-wavelength signals in a manner resembling colour opponency^10^. However, rod and cone signals are integrated at the level of the melanopsin-expressing retinal ganglion cells, which, as far as is currently known, provide the only retinal projection to the central clock. Furthermore, existing evidence suggests that this retinal integration does not itself exhibit the colour opponency observed within the SCN^15^. It therefore remains unclear whether the central clock receives an already separated colour signal from the retina that has yet to be identified, or whether the retina instead provides a more fixed, collapsed signal that must be unpacked within the clock itself. Bayesian models provide a way to make progress on this problem without first resolving the anatomy, because candidate rules for combining intensity and wavelength within the central clock make different predictions about the information that must be available in the retinal signal. The reliability-weighted integration proposed here, in which the balance between signals changes according to how informative each cue currently is, is analogous to optimal cue combination in other sensory systems^27^. This predicts a graded shift in cue dominance rather than an all-or-none switch, requiring access to the reliability of each cue. A fixed, collapsed retinal signal could not carry this information, implying that these computations must either be preserved or reconstructed somewhere downstream of the retina. The same logic, that the statistical signature of cue combination can reveal the existence of separate underlying estimates without directly observing them, has previously been used to demonstrate that the brain maintains independent estimates from different sensory modalities before combining them, such as vision and touch for spatial estimates^27^, and has been used to argue for multiple, modality-specific timing mechanisms rather than a single shared clock for sub-second time perception^44^. The same reasoning applies here. Demonstrating that the clock’s combined estimate follows the characteristic signature of reliability-weighted integration would itself provide evidence that intensity and wavelength are represented as separate estimates before being combined. A logical extension of this model, although outside the scope of this paper, is that reliability-weighted integration would allow non-photic cues to influence central clock function, for instance in populations without light input to the clock, such as the blind^45,46^. Indeed, it would be the exception rather than the rule if the central circadian clock did not integrate multiple sources of information, since this is a fundamental principle throughout mammalian neurobiology.

In our model, once intensity and wavelength are combined, the clock still needs to weigh that combined signal against its own internal estimate of astronomical time. We describe this as how much the clock “trusts” the incoming signal, shorthand for the width of the prior. A narrow prior means high confidence in the estimate of astronomical time and therefore resistance to resetting. A broad prior means high uncertainty and ready updating. The width of this prior should itself depend on how reliable the light-dark cycle has been in recent experience. A stable, regular cycle, where dawn and dusk recur at consistent times day after day, allows the clock to accumulate confidence that its internal estimate of astronomical time is accurate, narrowing the prior and making it more resistant to resetting by any single exposure. Conversely, an irregular or unpredictable light-dark cycle, or a long period without reliable light input at all, leaves the clock’s estimate poorly constrained, broadening the prior and increasing its sensitivity to whatever signal it does receive. This offers an account of why prior light history changes photic sensitivity: after prolonged exposure to constant darkness, rodents show phase shifts several-fold larger in response to the same light pulse than after only a single day of darkness^47^, and similar effects have been found in humans^35^, consistent with a prior that has broadened due to a prolonged absence of reliable timing information, rather than any change in the light-detection machinery itself.

A possible mechanism for computing prior uncertainty may lie in the internal synchrony of the clock’s neurons. Individual neurons within the SCN can act as independent timekeepers, displaying their own intrinsic circadian period^8^. Others have already proposed that internal synchrony among SCN neurons predicts phase resetting, although the evidence is mixed: a less internally synchronised clock can display greater phase resetting than a more synchronised one^48,49^, but the reverse has also been found^50^. Oscillator theory could potentially explain the first finding, as a population of weakly coupled oscillators has a lower amplitude and is more readily shifted by the same stimulus^51^, but it cannot account for the second. A Bayesian update can, because the size of the shift depends not only on the uncertainty of the prior, but also on the reliability of the light cue and the prediction it is compared against. Here we offer a computational reason why this might be functionally useful. If individual SCN neurons carry their own noisy estimate of astronomical time, their own individual “vote”, then the populations collective confidence is simply how much those votes agree: close agreement gives a narrow, confident prior, while disagreement gives a broad, uncertain one. This gives a testable prediction: the more desynchronised the population, the larger the phase shift it should show to the same light stimulus. However, an alternative account could argue that the phase shift is the light signal scaled by a synchrony-dependent gain factor, with a less synchronised network producing a bigger output to the same light. Such a gain curve can, in principle, be fit to any shape of the clock’s response to light, including at intermediate levels of synchrony, but it cannot constrain that shape in advance, the way a Bayesian prior-likelihood combination does: the relationship between prior width and the size of a phase update follows specifically from how prior and likelihood combine under Bayes’ rule. If light history and synchrony are genuinely independent components of the prior, and the likelihood is itself shaped by reliability weighting between cues, this points to something beyond parameter separability: different populations of SCN neurons would be required to serve different computational functions, some tracking the reliability of incoming cues to form a likelihood, others holding a running estimate of phase updated by that likelihood, with the confidence of that estimate read off from how well those neurons agree with one another. Research into the central clock is advancing, and it appears to be more complex than first thought. Whether it can hold computations consistent with Bayesian inference remains to be seen, but it appears plausible and opens a new avenue for circadian research.

## Notes

### Competing Interest Statement

The authors have declared no competing interest.

